# Belief updating in decision-variable space: past decisions with finer granularity attract future ones more strongly

**DOI:** 10.1101/2024.08.20.608767

**Authors:** Heeseung Lee, Jaeseob Lim, Sang-Hun Lee

## Abstract

When engaged in decision-making tasks, humans are known to create decision variables. Much effort has focused on the cognitive processes involved in forming decision variables. However, there is limited understanding of how decision variables, once formed, are utilized to adapt to the environment. We reason that decision-makers would benefit from updating the belief of decision-variable. As one such belief updating, we hypothesize that a decision commitment restricts the probabilistic belief distribution of decision variable to a range corresponding to that decision. This implies that past decisions not only attract future ones but also exert a greater pull when those decisions are made with finer granularity— dubbed ‘the granularity effect.’ Here, we present the findings of seven psychophysical experiments that confirm these implications. Further, as a unified account of the granularity effect, we offer a Bayesian model. Our work demonstrates how humans leverage the decision-variable to effectively adapt to their surroundings.

## INTRODUCTION

Making sound decisions is crucial for all living beings to survive and thrive, enabling them to take appropriate courses of action in diverse situations they face. All commanding decision theories^1-5^, despite their nuanced differences in formalism, point towards the need for constructing general cognitive quantity that can accommodate various sensory-motor mappings^6-8^. Such a need entails creating an internal space for abstract representation, where decision variables (DVs) are encoded through the integration of current evidence, prior knowledge, and learned cost^9,10^. Within this representational space^11^, dubbed ‘DV space,’ various operations are then carried out on DV to guide our course of action^2^. The DV space holds great significance in understanding the abstract nature and general applicability of adaptive intelligence^12^. The encoded state (value) of DV is *abstract* because it is not tied to the sensorimotor specifics of a task and thus *generalizable* because the DV formed for a task with particular sensorimotor specifics can be utilized to guide the performance in other tasks with different sensorimotor specifics, provided that the DV carries information shared across those tasks.

Due to its abstract and generalizable nature, the DV space has been considered a paragon of the cognitive mind’s ability to construct goal-directed mental playgrounds for intelligent behavior^13,14^. For this reason, extensive theoretical and empirical studies have been dedicated to elucidating the deliberate process of abstracting DVs from their sources and the commitment process of making a categorical choice^1,3,15-26^. These efforts substantially advanced our understanding of the deliberation and commitment processes of decision-making, including the origins of suboptimal DV encoding^27-29^ and the speed-accuracy tradeoff during DV formation^30-33^.

In stark contrast, much less effort has been put into understanding the cognitive processes that occur after DVs are encoded. There are good reasons to believe that these post-encoding processes are crucial in fostering adaptive behavior. Since only task-essential—but not inessential—elements are selectively integrated into DVs, DVs contain compact information about the otherwise almost infinitely complex environment^34^. In this sense, the DV space can be considered a complexity-reduced space where cognitive operations can be efficiently deployed to access the task-essential information with minimal computational cost^35^. In principle, since the DV space can embed continuous manifolds like other representational spaces (i.e., sensory and motor spaces), a specific probability distribution (belief) over DV states can be formed within the DV space. This raises the possibility that a belief about DV can be used to signify the degree of uncertainty, such as ‘decision confidence^36,37^,’ or updated over time to take advantage of stability in the environment^38^. Certainly, such computations are known to occur in the sensory and motor spaces. However, although this line of reasoning suggests that probabilistic computations are likely to be actively carried out in the DV space to steer adaptive behavior, this possibility has yet to be empirically validated.

As one potential adaptive operation in the DV space, we considered the cognitive act of committing to a decision with varying levels of granularity. A decision commitment with a specific level of granularity corresponds to confining the belief about the DV state within a specific region in the DV space. Suppose you are asked to sort the apples on a farm by size. To perform this task, you need to form a DV by calculating the size quantile of an apple: the probability that a specific apple is larger than any other apple on the farm. When asked to sort the apples into ‘small’ or ‘large’ classes, you are making decisions with a granularity of 2 (Figure 1A, top). Here, committing to the ‘small’ choice corresponds to restricting your current belief about the DV state within the lower half of the range (DV = [0, 1/2]; Figure 1B, top left). Importantly, the same commitment to the ‘small’ choice would have a different consequence when asked to perform a ternary decision-making task with a granularity of 3 (Figure 1A, bottom). Here, your belief about the DV state is restricted within the lowest third of the range (DV = [0, 1/3]; Figure 1B, bottom left). From this example, one can readily intuit that once a decision is committed, the width of DV belief narrows down linearly with decision granularity. Critically, if humans adaptively use their current belief as a prior expectation for future decision-making^39,40^ in the DV space—as they do in the sensory space^41-47^, the connection between decision granularity and belief width has a significant implication on temporal dependencies in decision-making^48^. Specifically, such belief propagation in the DV space implies that as the granularity in the current episode increases, the chosen DV state in that episode will exert a stronger attraction on the future episode’s DV state. This is because a narrower posterior in the current episode leads to a narrower prior in the next episode (Figure 1B), which more strongly attracts the likelihood function of DV. Consequently, as the current decision becomes more granular, it is increasingly likely to draw choices toward itself in the future (Figure 1C).

**Figure 1.**
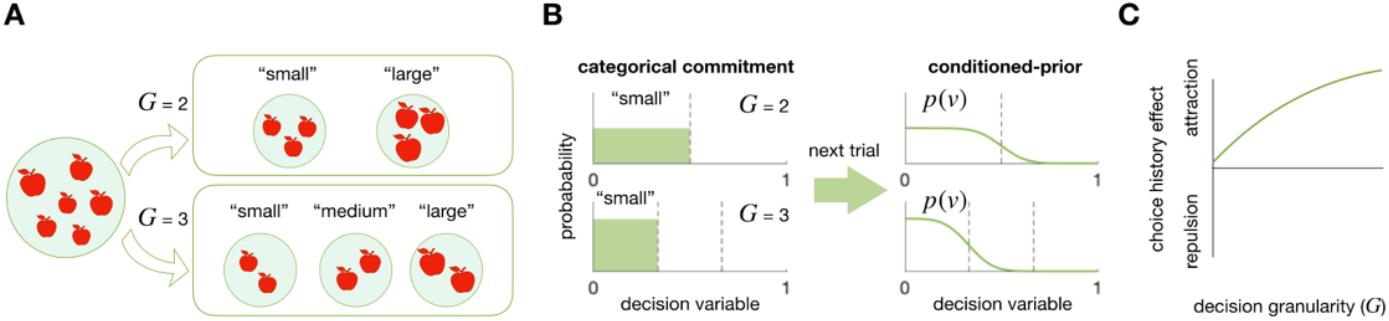
Decision granularity and its hypothetical impact on subsequent decision-making. (A) Decision commitment with varying levels of granularity (*G*). As the number of size classes into which apples are sorted increases, decision-makers are required to commit to a decision with a higher level of *G*. (B) Belief restriction by decision commitment (left) and prior updating in a subsequent trial (right) in the DV space. The higher the level of granularity, the more restricted and stronger the prior belief becomes. (C) Modulation of the choice-attraction bias by decision granularity. The granularity-dependent belief updating process depicted in panel B implies that the choice-attraction bias increases with the level of granularity.

To verify this implication, we crafted an experimental paradigm in which human individuals had to commit to a range of abstract values in the DV space with varying degrees of decision granularity. As implied, we found that the attractive bias toward the previous choice becomes stronger as decision granularity increases, a phenomenon we will call the “granularity effect.” Reflecting the *abstract* nature of DVs, the granularity effect was unaffected by the modality of sensory input and the arrangement of motor output, consistently observed regardless of whether the stimuli were visual or auditory and whether the choice-to-action mapping was rearranged. Reflecting the *generalizable* nature of DVs, the granularity effect was observed even when decision commitment was followed by different sensorimotor specifics, such as point estimation, as long as the following task needs to be performed in the same DV space.

Furthermore, reflecting the *separation* of the DV space from the sensory space, the granularity effect was no longer observed when decision commitment was followed by a task that only required accessing the sensory space alone, but not the DV space. Lastly, by building a Bayesian network model^39,46,49,50^, we offer a principled account of the observed granularity effect by demonstrating that it can arise from the adaptive belief propagation over sequential episodes in the abstract and generalizable DV space, separated from the stimulus and action spaces.

Our findings reveal a previously unrecognized adaptive process in the DV space. This highlights the human intellect’s adeptness in adapting probabilistic belief distributions to recent experiences within the abstract and generalizable DV space, where task-essential information is represented concisely, invariant to sensory-motor specifics.

## RESULTS

### Task paradigm and rationale to probe the granularity effect

To probe the granularity effect in the DV space, we asked participants to perform magnitude classification tasks^46,49-51^ where the number of classes varied (2, 4, or 8) either block-wise or trial-wise. In each trial, participants viewed a stimulus and sorted its magnitude (e.g., size or pitch) into one of the pre-designated classes (Figure 2A). Across trials, magnitudes were randomly sampled from a single fixed distribution for a given type of magnitude. By doing so, the stimulus factor (*S*) and the granularity factor (*G*) were orthogonally manipulated.

**Figure 2.**
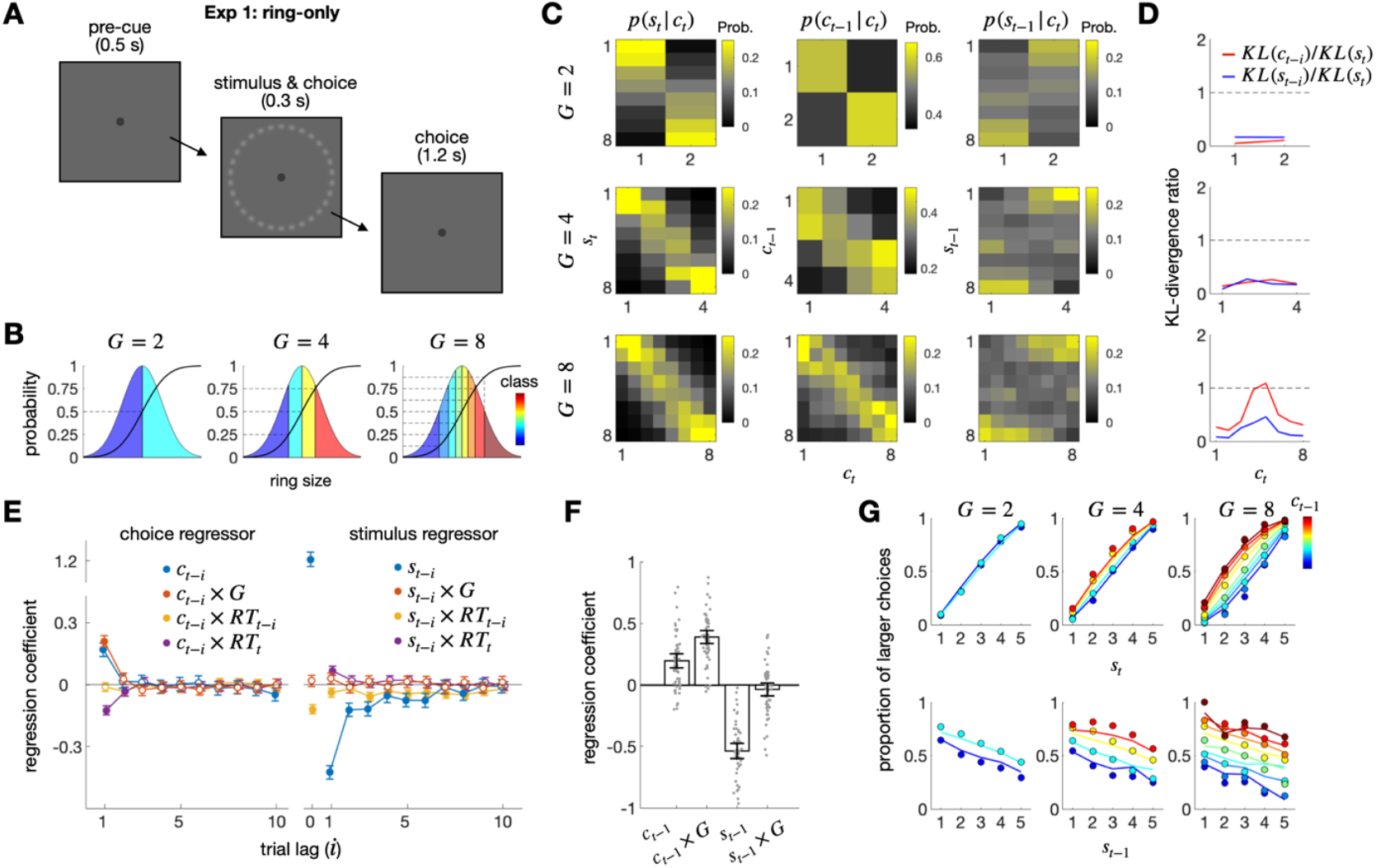
Exp 1: Ring-only experiment. (A) Schematic illustrations of the ring size classification task. After viewing a ring, participants classified its size into one of multiple classes. (B) Definition of classes with varying degrees of granularity (*G*). A continuous normal probability density function is divided into *G* parts of equal proportion, as indicated by the cumulative density function. The class of a stimulus corresponds to the order of the part to which the stimulus belongs. (C) Conditional probability distributions of the current stimulus (*s*_*t*_), previous choice (*c*_*t*−1_), and previous stimulus (*s*_*t*−1_) given the current choice (*c*_*t*_). Each column of the panels displays the probability distribution of *s*_*t*_ (left), *c*_*t*−1_ (middle), or *s*_*t*−1_ (right). Each row of the panels shows a different granularity condition. Continuous values of stimulus size are binned into eight bins. (D) The influences of the previous choice and stimulus on each state of the current choice, relative to the influence of the current stimulus, captured by Kullback-Leibler Divergence (KL) measures. Each KL measure (*KL*(*s*_*t*_), *KL*(*c*_*t*−1_), and *KL*(*s*_*t*−1_)) quantifies the degree to which the corresponding conditional distribution in panel C deviates from the uniform distribution. (E) Coefficients of the regression model fitted to the pooled choices of all participants for the regressors involving the previous choice (left) and the previous stimulus (right) in the size classification task. *x* × *y* denotes the interaction term between variables *x* and *y*. Here and thereafter, unless specified otherwise, the confidence interval denotes a 95% confidence interval, and the filled circles indicate the coefficients that significantly deviate from 0 after being controlled for FDR. (F) Coefficients of the regression model fitted to the choices of each participant (gray dots). Vertical bars indicate the mean of the coefficients across participants. (G) The granularity effect visualized in psychometric curves. The proportion of larger choices, which belongs to an upper half of the classes, conditioned on *s*_*t*_ and *c*_*t*−1_ (upper) or *s*_*t*−1_ and *c*_*t*−1_ (lower) are shown for each granularity condition. The color hues of the symbols and lines indicate the class of the previous choice. The circles and lines represent human choices and the predictions of the regression model in panel E, respectively.

We assume that decision-makers carry out the magnitude classification task using the following algorithm: (i) they embed a line manifold of relative magnitude^52^—a relative position within a population of interest—within the DV space, and (ii) they commit to a specific class by picking a subsection of the manifold where the item of interest falls. We then hypothesize that an act of making a decision commits the decision-maker to a specific belief about the DV states. This limits the probability distribution to the subsection of the manifold linked to the chosen class within the DV space (Figure 1B, left). We further hypothesize that this *limited* posterior belief in the current trial propagates into the next trial, serving as a prior belief about the DV states (Figure 1B, right) by pulling the likelihood belief given the sensory evidence in that trial, leading to the attraction bias in choice history effect: currently chosen classes are likely to be similar to previously chosen classes. We will refer to this hypothesis as the ‘belief-updating-in-DV’ hypothesis. Critically, the belief-updating-in-DV hypothesis predicts that this choice-attraction bias becomes stronger as granularity increases—the granularity effect (Figure 1C). This is because decision commitments with finer granularity result in narrower priors, which in turn exert stronger influences on the next DV formation.

With the paradigm described above, we probed the influences of two history factors—the previous stimulus (*s*_*t*−*i*_) and the previous choice (*c*_*t*−*i*_) —on the current choice (*c*_*t*_) and assessed the modulatory influence of decision granularity (*G*) on these two history effects under diverse experimental settings. In doing so, we will focus on testing the implications of the belief-updating-in-DV hypothesis: (i) the current choice is attracted toward the previous choice (choice-attraction bias); (ii) the strength of this attraction increases as a function of decision granularity (granularity effect). Once we establish these two implications, we will investigate a range of additional nuanced implications of the hypothesis: whether the presence or absence of the granularity effect aligns with the essential characteristics of the DV space—being abstract, generalizable, and distinct from sensory and motor processes spaces.

### Exp 1: The modulation of the choice-attraction bias by granularity in ring size stimuli

We began by confirming the choice-attraction bias and its modulation by decision granularity in the visual domain for an adequate number of participants. In the first experiment (Exp 1), fifty-eight participants were tasked to sort a sequence of ring stimuli by size into discrete classes, a method proven effective in studying history effects in our previous research (Figure 2A,B)^46,49-51^.

Initially, we calculated the probability distribution of the previous choice’s states, conditioned on the current choice’s states (*p*(*c*_*t*−1_|*c*_*t*_ = *i*); Figure 2C, each column of the center panels). The higher densities of the previously chosen states near the value corresponding to the currently chosen state (as indicated by the bright yellow squares surrounding the diagonal of the center panels in Figure 2C) visually capture the attractive influence of the previous choice on the current choice (‘choice-attraction bias’). On the contrary, opposite patterns were observed when the probability distribution of the previous stimulus’s states was conditioned on any given state *k* (i.e., class of size) of the current choice (*p*(*s*_*t*−1_|*c*_*t*_ = *k*); Figure 2C, each column of the right panels), indicating the previous stimulus’s repulsive influence on the current choice (‘stimulus-repulsion bias’).

To compare the strength of the choice-attraction and stimulus-repulsion biases across granularity levels, we (i) quantified the extent to which the distributions deviate from a uniform distribution, which corresponds to ‘no influence,’ using Kullback-Leibler divergence (*KL*) and then (ii) normalized these quantities by dividing them by the *KL* quantities for the probability distribution of the current stimulus’s states conditioned on the corresponding state of the current choice (Figure 2C, each column of the left panels; see STAR methods for details). Thus, these normalized *KL* values (*KL*(*c*_*t*−1_|*c*_*t*_ = *k*)/ *KL*(*s*_*t*_|*c*_*t*_ = *k*) and *KL*(*s*_*t*−1_|*c*_*t*_ = *k*)/*KL*(*s*_*t*_|*c*_*t*_ = *k*); Figure 2D) signify the strength of the previous choice and stimulus on the current choice, compared to the influence of the current stimulus on the current choice: ‘0’ indicates no influence, while ‘1’ indicates an influence equal to that of the current stimulus. We found that as decision granularity increased, the normalized *KL* values substantially increased for the previous choice (the red lines in Figure 2D). We also note that the granularity effect on the choice-attraction bias, although consistently observed across all the states of the current choice, was most pronounced for the intermediate states.

While the choice-conditioned probability distributions effectively visualize the historical effects, this method is limited as a rigorous statistical test because other variables, such as previous stimuli and choices further back than one trial lag, as well as response times (RTs), should be concurrently conditioned to isolate the unique contributions of regressors. In principle, it allows for the possibility that the observed effect may not arise from the factor of interest, but instead from associated “lurking factors”^53^. To address this limitation, we conducted a multiple regression analysis in which the current choice was simultaneously regressed onto the previous choice and stimulus while including decision granularity (*G*) and RTs as modulatory factors. Choices farther than the 1 trial lag (*c*_*t*−*i*_, where *i* > 1) are included because the history effect can exert more than the adjacent trials. The previous stimulus (*s*_*t*−*i*_) was included because it is correlated with the previous choice and tends to repel the current choice^46,54-56^, as observed above. In addition, the interaction terms involving response time (*c*_*t*−*i*_ × *RT*_*t*_; *c*_*t*−*i*_ × *RT*_*t*−*i*_) were included because RTs are correlated with task difficulty, which in turn is likely to covary with decision granularity and can modulate the attraction effect of the previous choice on the current choice^56-59^.

After defining the co-regressors, we added several features to the multiple regression model to fairly compare the history effects across varying levels of decision granularity. First, as a regressand, the current choice was treated as binary without considering decision granularity (*c*_*t*_ ∈ [0,1]): ‘0’ and ‘1’ correspond to the lower and upper halves of the current choice’s states, respectively. Second, as a regressor, the previous choice was decomposed into three orthogonally ordered binary codes 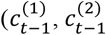, and 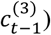. Since each order of the binary code carries one bit of information, decision granularities of 2, 4, and 8 can be sufficiently and adequately captured by the first-, second-, and third-order codes, respectively. Specifically, the first-order component captures the influence of the binarized states of the previous choice 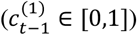: ‘0’ and ‘1’ correspond to the lower and upper halves of the previous choice’s states, respectively. The second component captures the influences of the second-order binary states of the previous choice with a decision granularity of 4 or 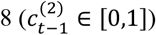: ‘0’ and ‘1’ correspond to the lower and upper halves of the states *within* each binarized state of the first-order component. Likewise, the third component captures the influences of the third-order binary states of the previous choice with a decision granularity of 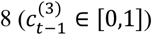: ‘0’ and ‘1’ correspond to the lower and upper halves of the states *within* each binarized state of the second-order component (see Table 1 in STAR methods for details). This decomposition enables the regression model to fairly compare the influence of the ‘binarized’ previous choice at varying levels of granularity while thoroughly evaluating the finer, higher-order influences of the previous choice. As mentioned earlier, ensuring ‘exhaustive regression’ is important; otherwise, the unexplained higher-order influences of the previous choice could be mistakenly attributed to the previous stimulus due to the strong connection between stimuli and choices. We emphasize that although the higher-order choice regressors were included in the model, only the coefficients of the first-order choice regressor were used to compare the effects of choice history across the three granularity conditions. Third, the combined influence of all the regressors was linked to the regressand through the probit function. This multiple regression model (Equation 1), which includes the three features mentioned above, will henceforth be referred to as the ‘decomposition model.’

**Table 1.** The decomposition of choice. The decomposition encodes each choice (*c*_*t*_) into three values 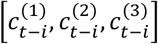 for each granularity, respectively.

| $c_t$ | $G_t = 2$ | $G_t = 4$ | $G_t = 8$ |
| --- | --- | --- | --- |
| 1 | [0,0,0] | [0,0,0] | [0,0,0] |
| 2 | [1,0,0] | [0,1,0] | [0,0,1] |
| 3 | - | [1,0,0] | [0,1,0] |
| 4 | - | [1,1,0] | [0,1,1] |
| 5 | - | - | [1,0,0] |
| 6 | - | - | [1,0,1] |
| 7 | - | - | [1,1,0] |
| 8 | - | - | [1,1,1] |

By fitting the coefficients of the decomposition model to the data from Exp 1, at both group and individual levels, we evaluated the significance and strength of the history effects. The group-level results confirmed the history effects observed through the probability distributions conditioned on the previous choice and stimulus: the previous choice and stimulus attracted (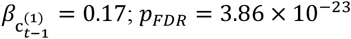; ‘choice-attraction bias’) and repelled (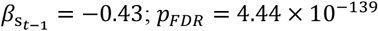; ‘stimulus-repulsion bias’), respectively, the current choice (blue dots in Figure 2E), as reported previously^42,46,56^. In addition, the choice-attraction bias became stronger as response time became shorter (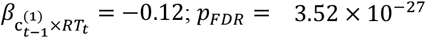; purple dots in Figure 2E left), as reported by previous research^58^. Importantly, the granularity effect, a phenomenon of interest and novelty, was significant for the previous choice: as granularity increased, the choice-attraction bias increased (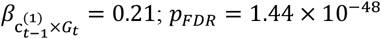; the leftmost solid red dot in Figure 2E left). In contrast, such a modulatory effect of granularity was absent for the stimulus-repulsion bias (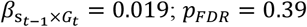; the leftmost empty red dots in Figure 2E right). The individual-level results (Figure 2F) corroborated the group-level findings by showing a significant choice-attraction bias 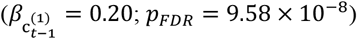, a significant stimulus-repulsion bias 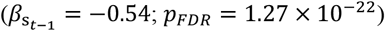, and a significant granularity effect only for the previous choice 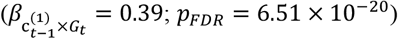 but not for the previous stimulus 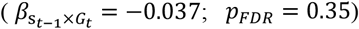. This selective modulation of the choice attraction bias by granularity is consistent with our hypothesis in that the belief propagation constrained by granularity occurs in the DV space, which is *separate from* the sensory (stimulus representation) space.

To visually summarize the history effects captured by the multiple regression, we plotted the psychometric curves as functions of the current stimulus (*ψ*(*s*_*t*_); Figure 2G, top) and the previous stimulus (*ψ*(*s*_*t*−1_); Figure 2G, bottom), each conditioned on the state of the previous choice and the levels of granularity. The decomposition model accurately predicted the observed proportions of the human participants’ binarized ‘large’ choices (as indicated by the close correspondence between the lines and circles in Figure 2G; *R*^2^ = 0.45 for group-level; *R*^2^ = 0.59 ± 0.017 for individual-level, mean±95%CI). *ψ*(*s*_*t*_) was elevated upward as the previous choice’s state became ‘*larger*’ (labeled with colors in the top panels of Figure 2G), indicating the presence of the choice-attraction bias, with the degree of elevation increasing as a function of decision granularity (left to right columns in the top panels of Figure 2G top). The negative slope of *ψ*(*s*_*t*−1_), which indicates the presence of the stimulus-repulsion bias, remained constant across the three levels of granularity, indicating the lack of granularity effect for the previous stimulus. In contrast, *ψ*(*s*_*t*−1_) was elevated upward as the previous choice’s state became ‘*larger*’ (labeled with colors in the bottom panels of Figure 2G), once again indicating the presence of the choice-attraction bias. Likewise, the degree of elevation increased as a function of decision granularity (left to right columns in the bottom panels of Figure 2G top), confirming the granularity effect on the choice-attraction bias.

### Exp 2: The modulation of the choice-attraction bias by granularity in sound pitch stimuli

An important characteristic of the DV space is its independence from the sensory sources that form DVs—*abstractness*. This implies that the granularity effect should occur irrespective of the specifics of sensory modality, as long as decision commitment takes place in the DV space. To verify this implication, we conducted the second experiment (Exp 2), in which the same fifty-eight individuals who participated in Exp 1 performed a sound pitch classification task (Figure 3A,B). This task had the same structure as the ring size classification task, except that auditory stimuli were sorted by their pitch.

**Figure 3.**
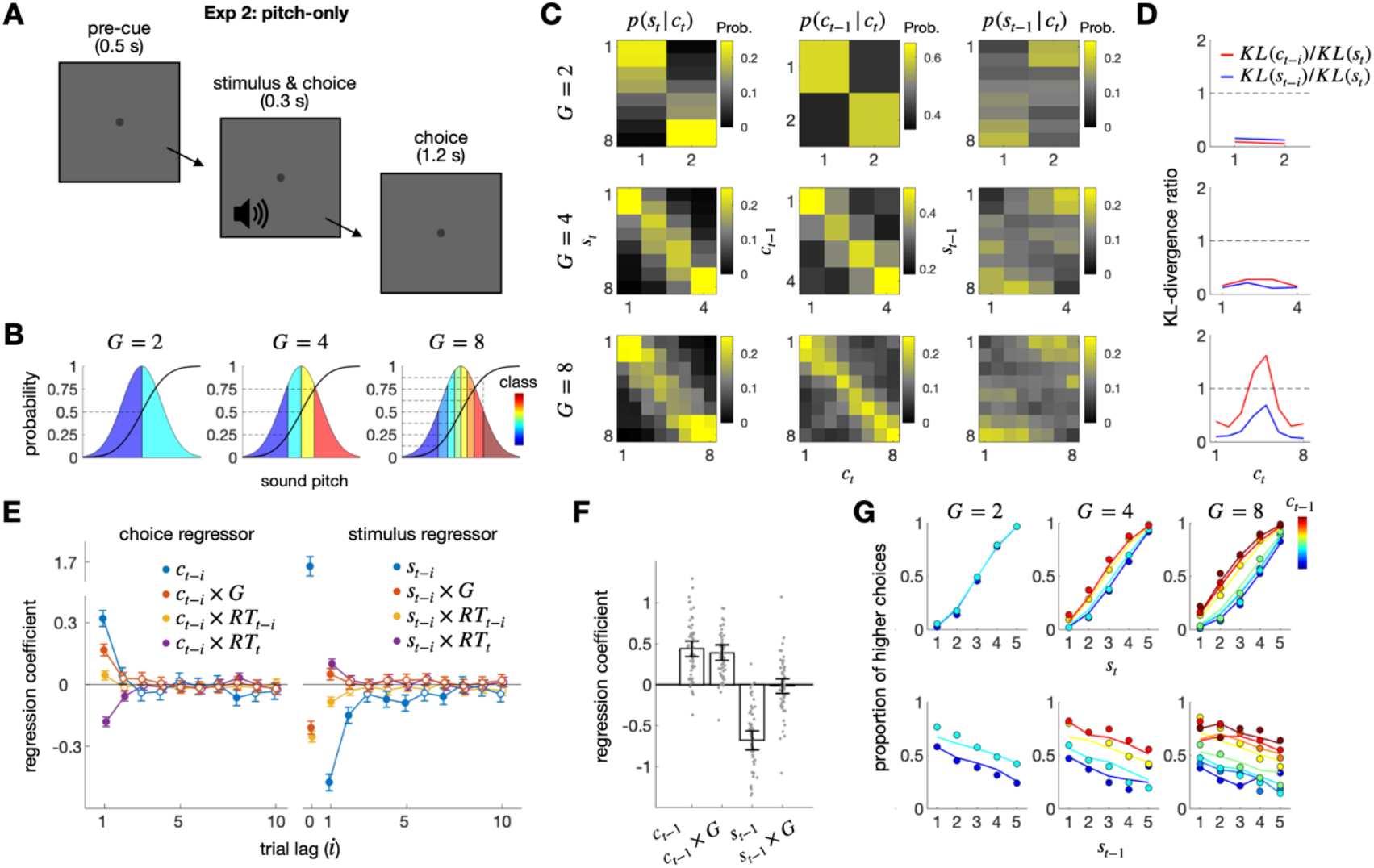
Exp 2: Pitch-only experiment. (A) Schematic illustrations of the sound pitch classification task. After hearing a beep sound, participants classified its pitch into one of multiple classes. (B) Definition of classes with varying degrees of granularity (*G*). (C) Conditional probability distributions of the current stimulus (*s*_*t*_), previous choice (*c*_*t*−1_), and previous stimulus (*s*_*t*−1_) given the current choice (*c*_*t*_). (D) The influences of the previous choice and stimulus on each state of the current choice, relative to the influence of the current stimulus, captured by Kullback-Leibler Divergence (KL) measures. (E) Coefficients of the regression model fitted to the pooled choices of all participants for the regressors involving the previous choice (left) and the previous stimulus (right) in the pitch classification task. (F) Coefficients of the regression model fitted to the choices of each participant (gray dots) (G) The granularity effect visualized in psychometric curves. See the captions of Figure 2 for details, as the format matches that of Figure 2.

The same set of analyses applied to Exp 1 was used to analyze the data from Exp 2. Results from Exp 2 supported those from Exp 1, consistently demonstrating the choice-attraction bias, the stimulus-repulsion bias, and the granularity effect for the choice-attraction bias in the choice-conditioned distributions, the multiple regression analyses at both group and individual levels, and the psychometric curves.

The probability distributions of the states of the previous choice and stimulus conditioned on the current choice’s states were similar to those in Exp 2 (see Figure 3C, D compared to Figure 2C,D), clearly indicating that the choice-attraction bias increased in strength as decision granularity increased. When the decomposition model was fitted to the data at the group level, its coefficients were similar to the corresponding values in the size task (see Figure 3E compared to Figure 2E), exhibiting the choice-attraction bias 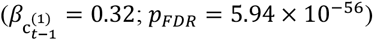, the stimulus-repulsion bias 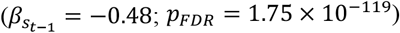, and the granularity effect for the choice-attraction bias 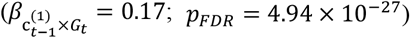. These results were also confirmed on an individual basis (Figure 3F; 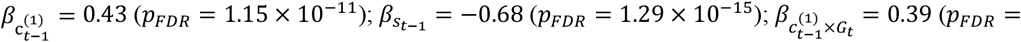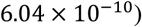). The stimulus-repulsion bias was weakly modulated by decision granularity at the group level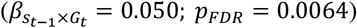, but it was not significant at the individual level 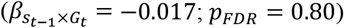. Once again, the decomposition model accurately captured the psychometric curves, accounting for the elevation of the curves due to the previous choice’s state as well as its modulation by decision granularity (Figure 3G).

### Exp 3: Source-specific choice-attraction bias and its modulation by granularity

We account for the choice-attraction bias based on belief updating in the DV space. This account has a rational basis: it leverages the statistical tendency that magnitudes sampled closely in time from the same population are similar to one another in the natural environment^38^. For instance, the temperatures show greater similarity from one day to the next than from one week to the next. Importantly, this environmental stability holds *true only when magnitudes are sampled from the same population*. There is no reason to believe that a magnitude sampled at time *t* from population A is similar to a magnitude sampled at time *t* + 1 from population B, as long as there is no relationship between the two populations.

For instance, as we walk in the woods, the relative pitch of bird chirping sounds I hear now does not provide any indication of the size of rocks I will encounter next. This implies that there should be no choice-attraction bias and, therefore, no granularity effect between choices of different magnitude populations if the bias reflects the brain’s rational strategy to exploit environmental stability.

We verified this implication in the third experiment (Exp 3), where the size and pitch tasks were randomly intermixed across trials (Figure 4A). When the magnitude source for classification was the same between consecutive trials, the history effects observed in Exp 1 and 2 were observed (Figure 4B,C): the current choice was attracted to the previous choice 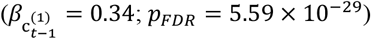, and repelled from the previous stimulus 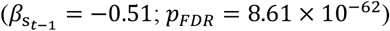; the choice-attraction bias was greater for a granularity of 4 than for a granularity of 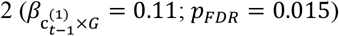. In contrast, when the magnitude source for classification differed between consecutive trials, none of the history effects observed in Exp 1 and 2 were observed (Figure 4C,D): neither the choice-attraction bias 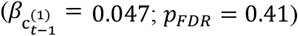, the stimulus-repulsion bias 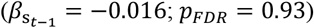, nor the granularity effect for the choice-attraction bias was observed 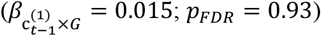.

**Figure 4.**
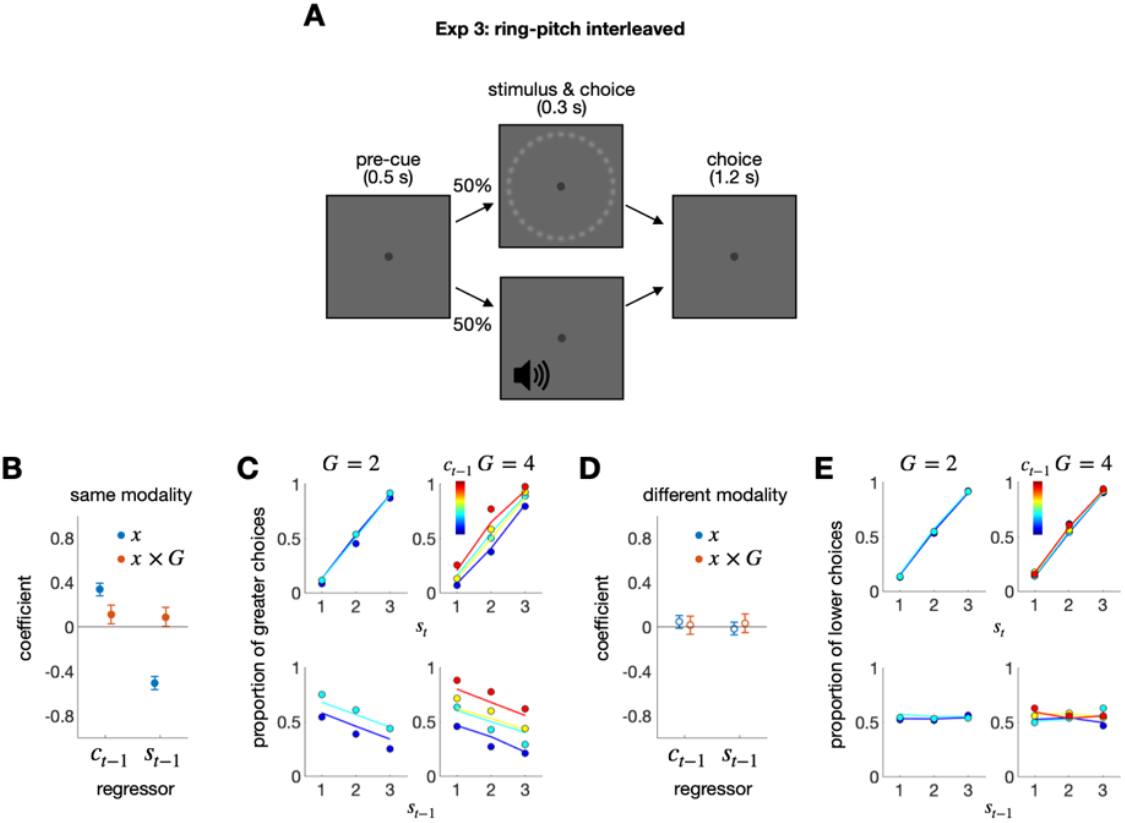
Exp 3: Ring-pitch interleaved experiment. (A) Schematic illustrations of the ring-pitch interleaved classification task. In each trial, a ring or pitch stimulus was randomly presented, and participants classified the stimulus into one of multiple classes. (B,D) Regression coefficients for the previous stimulus (*s*_*t*−1_) and choice (*c*_*t*−1_) factors and their interaction terms with the granularity factor (*s*_*t*−1_ × *G*; *c*_*t*−1_ × *G*), when the consecutive stimulus modalities were the same (B) or different (D). The filled circles indicate the coefficients that significantly deviate from 0 after control for FDR. (C,E) The psychometric curves for consecutive stimulus modalities being the same (C) or different (E). The proportion of ‘greater’ choices, which belong to the upper half of the classes, was conditioned on *s*_*t*_ and *c*_*t*−1_ (upper) or *s*_*t*−1_ and *c*_*t*−1_ (lower) for each granularity condition. The color hues of the symbols and lines indicate the class of the previous choice. The circles and lines represent human choices and the predictions of the regression model, respectively.

These results suggest that belief updating within the DV space occurs only when DVs are derived from the same source, specifically, the same distribution of task-relevant properties.

### Exp 4: Trial-to-trial prospective modulation of the choice-attraction bias by granularity

Until now, the granularity effect has only been demonstrated under experimental conditions where granularity varies block-wise but remains fixed trial-wise. In the fourth experiment (Exp 4), granularity was varied on a trial-to-trial basis because manipulating granularity trial-to-trial allows for putting our account of the granularity effect to a rigorous test by validating its prediction regarding the temporal direction of the effect (Figure 5A). According to our account, the extent to which the current choice is attracted toward the previous choice is determined by the level of belief granularity in the previous trial but not in the current trial. This implies that the current choice is regressed significantly onto the previous choice × previous granularity term 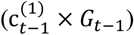 but not significantly onto the previous choice × current granularity term 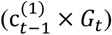.

**Figure 5.**
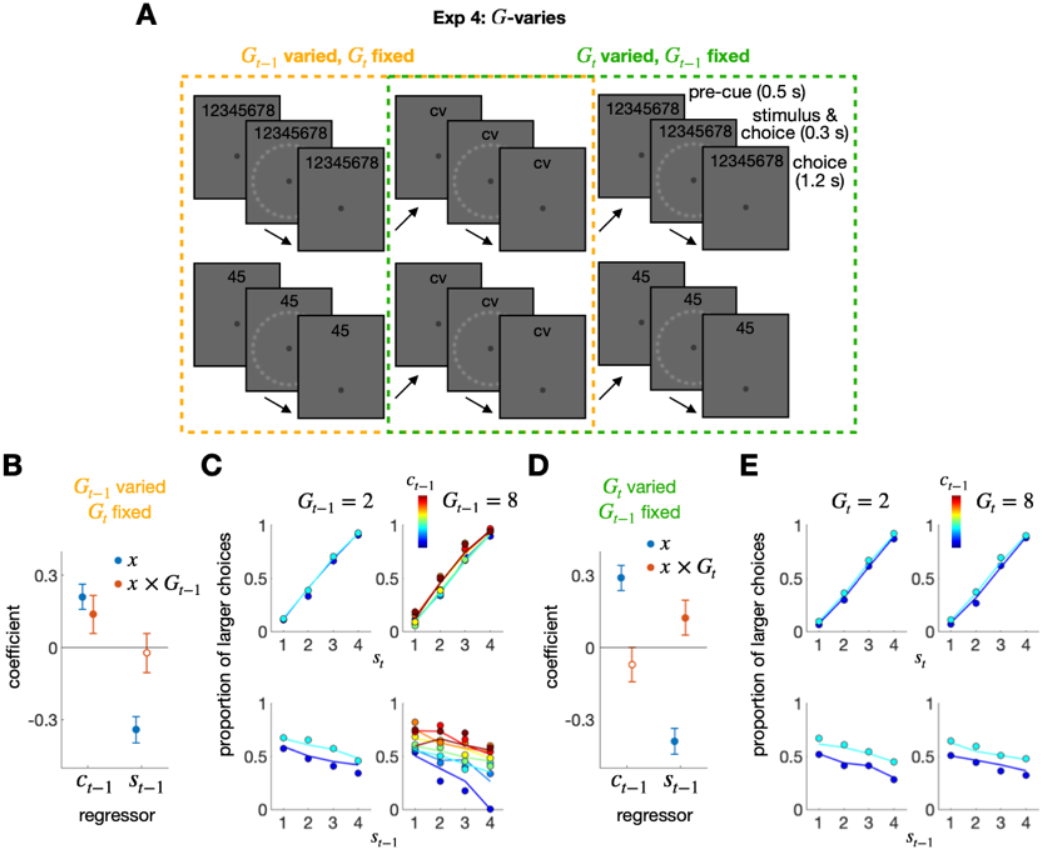
Exp 4: Granularity-varying experiment. (A) Experimental design. Exp 4 involved two types of blocks: one in which the level of granularity switched between 2 and 8 from trial to trial (top panels), and the other in which the level of granularity remained constant at 2 across trials (bottom panels). The decision granularity and response keys to be used were indicated by a string of numbers or letters shown at the top of the display. By sorting trials from the two types of blocks, the history effects were compared between two conditions: one in which the level of granularity was varied in the previous trial and fixed in the current trial (orange dashed box), and the other in which the level of granularity was fixed in the previous trial and varied in the current trial (green dashed box). (B,D) Regression coefficients for the previous stimulus (*s*_*t*−1_) and choice (*c*_*t*−1_) factors and their interaction terms with the granularity factor (*s*_*t*−1_ × *G*; *c*_*t*−1_ × *G*), when decision granularity was varied in the previous (B) or current (D) trial. The filled circles indicate the coefficients that significantly deviate from 0 after control for FDR. (C,E) The psychometric curves for decision granularity being varied in the previous (C) or current (E) trial. The proportion of ‘greater’ choices, which belong to the upper half of the classes, was conditioned on *s*_*t*_ and *c*_*t*−1_ (upper) or *s*_*t*−1_ and *c*_*t*−1_ (lower) for each granularity condition. The color hues of the symbols and lines indicate the class of the previous choice. The circles and lines represent human choices and the predictions of the regression model, respectively.

The results confirmed this implication: the choice-attraction bias increased as the previous granularity increased, with a fixed current granularity 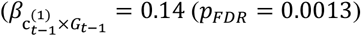; Figure 5B, C), but did not increase as the current granularity increased, with a fixed previous granularity 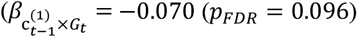; Figure 5D,E). These results indicate that the factor modulating the choice-attraction bias is not the decision granularity in the current trial, but rather in the previous trial.

### Exp 5, 6: Generalizability and separability of the granularity effect

The belief formed in a given DV space is *generalizable* within that space but *separable* from the sensory space where stimulus features are represented. In the current context, *generalizability* implies that the belief granularized through decision commitment should continue to influence the belief formed in the next trial, even if the sensorimotor specifics in the next trial are different from those in the previous trial, as long as the next task is carried out in the same DV space as the previous task. On the other hand, *separability* implies that the granularized belief in the previous trial should NOT influence the belief in the next trial, even if the same type of sensory stimuli is used in the following trial, as long as the next task is carried out in the sensory space rather than in the DV space. We tested these two implications in the fifth (Exp 5) and sixth (Exp 6) experiments.

Specifically, the size classification task was followed randomly by either a size scaling task or a size reproduction task (Figure 6A). Size magnitudes were randomly sampled from the fixed distribution, as before, while the number of classes varied between two and eight in the classification task to manipulate granularity. In the scaling task, observers estimated the relative size on a scale from 0 to 1. This task requires accessing the same DV space used in the classification task, where DVs are abstracted from placing perceived sizes within the overall population size distribution. In contrast, the reproduction task, in which observers reproduce the absolute size of the ring, does not require accessing the DV space at all; instead, it can be carried out by accessing the value encoded in the sensory space. This contrast in task structure predicts that the granularity in the previous classification-task trial selectively affects the choice-attraction bias in the subsequent trial of the scaling task, but not in the reproduction task.

**Figure 6.**
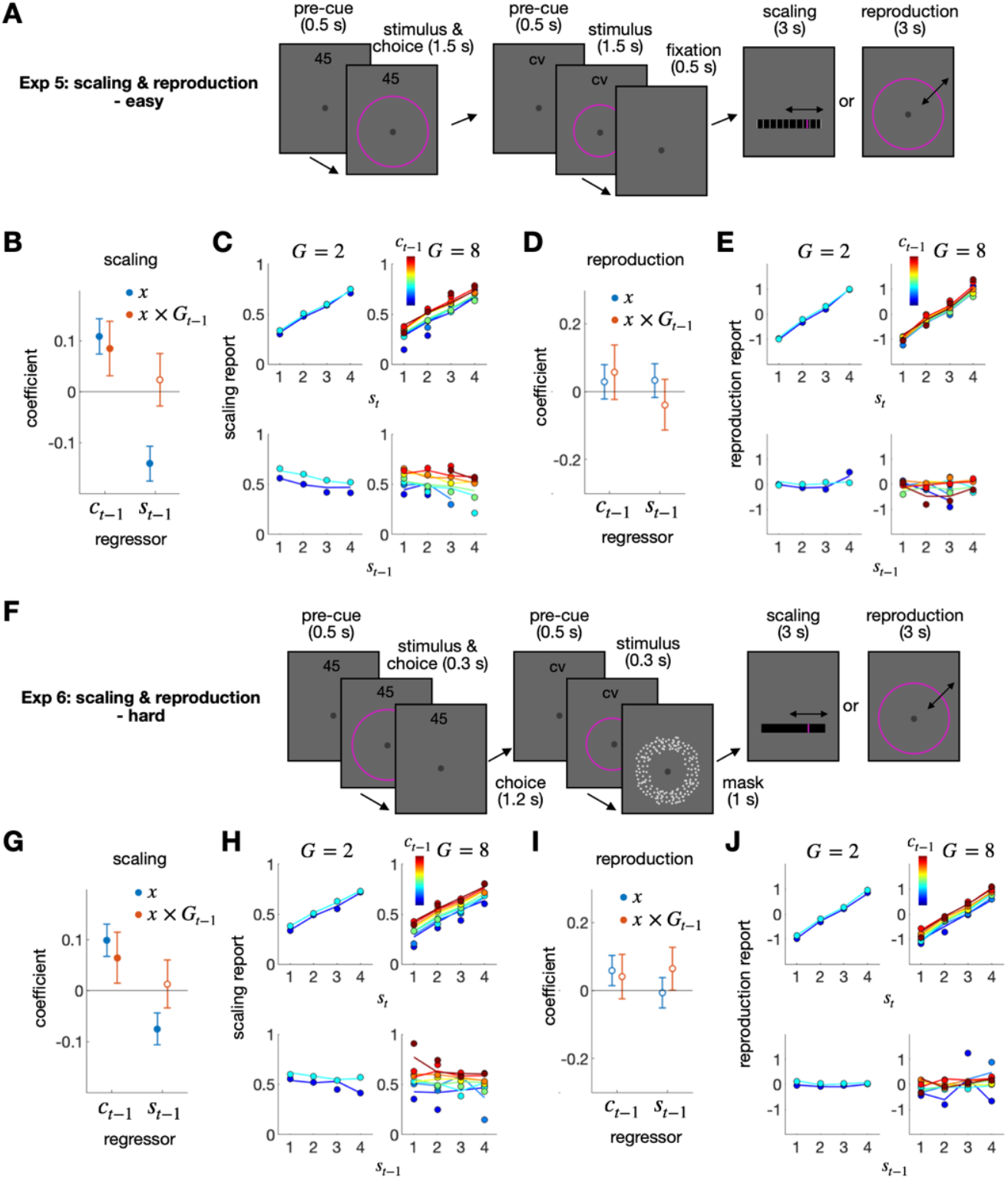
Exp 5 and 6: Scaling and reproduction after classification. (A) Experimental design for Exp 5. There were two types of blocks: the classification-task trial was followed either by the scaling-task trial or by the reproduction-task trial. The level of granularity was varied between 2 and 8 for the classification task. (B-E) Regression and psychometric curve results, shown separately for the two block types described in panel A. B,D, Regression coefficients for the previous stimulus (*s*_*t*−1_) and choice (*c*_*t*−1_) factors and their interaction terms with the granularity factor (*s*_*t*−1_ × *G*; *c*_*t*−1_ × *G*) are shown for the classification-followed-by-scalin-task trials (B) and for the classification-followed-by-reproduction-task trials (D). The filled circles indicate the coefficients that significantly deviate from 0 after control for FDR. (C,E) The scaling and the reproduction responses are conditioned on *s*_*t*_ and *c*_*t*−1_ (upper) or *s*_*t*−1_ and *c*_*t*−1_ (lower) for each granularity on the previous trial. The color hues of the symbols and lines indicate the class of the previous choice. The circles and lines represent human responses and the predictions of the regression model, respectively. (F) Experimental design for Exp 6. Exp 6 had the same design as Exp 5, except for differences in stimulus duration, the presence of a backward mask, and the inclusion of scaling ticks, which make Exp 6 more challenging than Exp 5. (G-J) Regression and psychometric curve results, shown separately for the two block types described in panel F. The figure format is consistent with panels B-E.

The prediction was confirmed: the scaling estimates were attracted toward the previous classification choice 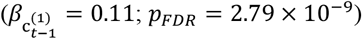, with the degree of attraction increasing as granularity increased 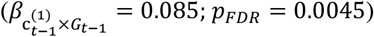 (Figure 6B,C), whereas the reproduction estimates were not influenced by the classification choice at all 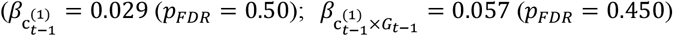; Figure 6D,E). This marked difference between the two tasks was robust: it held true when stimuli were shown briefly and followed by a backward mask (Figure 6F-J; scaling task: 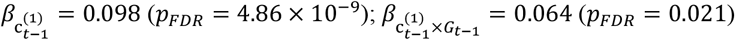; reproduction task: 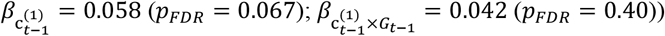.

These results suggest that the act of decision commitment narrows down the range of belief about the DV states and that this reduction has a generalizable impact on tasks that access the same DV space with different sensorimotor specifics but does not affect tasks that access only the sensory space, which is separate from the DV space.

### Exp 7: Action-space-invariant modulation of the choice-attraction bias by granularity

For two reasons, one might consider the action space as an alternative to the DV space for the origin of the granularity effect. First, the number of available actions increases as granularity increases (Figure 7A). This transformation of the action space, often called ‘action space shaping,’ is known to affect how agents learn a task, similar to reward shaping^60^. Second, in Exp 1, 2, 4, 5, and 6, committing to a specific choice perfectly aligns with taking action at a specific location in the action space for a given level of granularity (Figure 7A). This strong association between the action and DV spaces can obscure the source of the granularity effect. It should be taken seriously because the location of the response hand and the semantic magnitude are known to be automatically associated^61-64^. Additionally, the granularity effect was not observed in the different modality condition of Exp 3, where the DV space happened to be not aligned with the action space (Figure 7B). Therefore, we predicted that if the granularity effect originates from the action space, the current choice would be repelled from the previous choice by reversing the action space from the DV space arrangement trial-to-trial. Conversely, if the DV space induces the granularity effect, it would still be observed even when the action space is reversed, as long as the DV space remains unchanged aligned.

**Figure 7.**
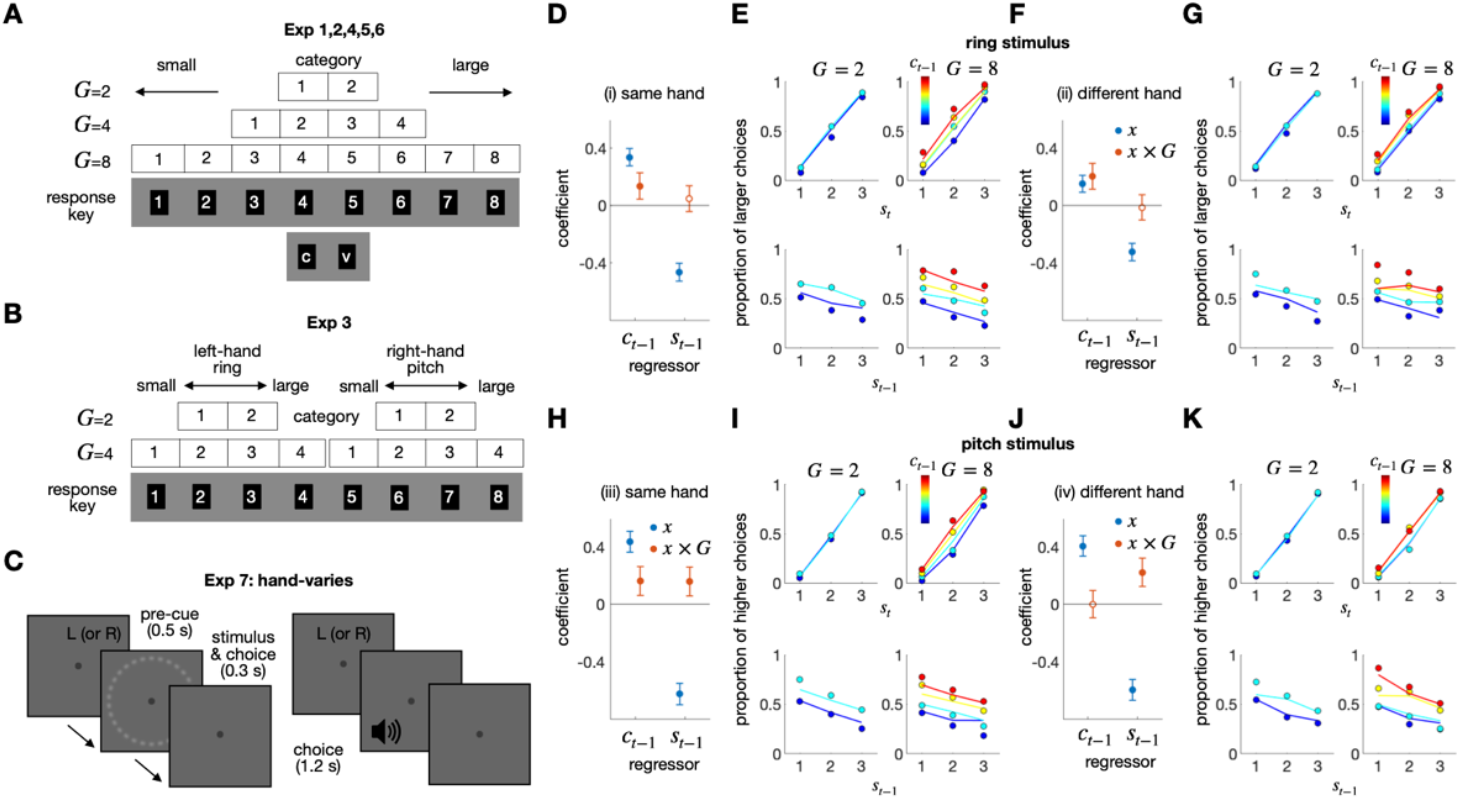
Results of Exp 7. (A,B) Spatial arrangement of response keys for Exp 1-6. A, Linear association between the class states and the locations in action space in Exp 1, 2, 4, 5, and 6. Within these experiments, the class identities were orderly mapped onto a fixed set of response keys arranged on the computer keyboard, which served as a constant action space. B, Nonlinear association between the class states and the locations in action space in Exp 3. Within each task type, the class identities were orderly mapped onto the fixed action space, while the action space differed between the two task types since the left and right hand were used for the ring size classification (left) and sound pitch classification (right) tasks, respectively. (C) Experimental design of Exp 7. Participants performed the ring size classification task and the sound pitch classification task in different blocks. Hand assignment was randomized from trial to trial. Participants were instructed which hand to use for a response by the letter displayed at the top of the screen at the beginning of each trial. There were four types of two consecutive trials: (i) ‘same-hand ring-size-classification’ trials, (ii) ‘different-hand ring-size-classification’ trials, (iii) ‘same-hand sound-pitch-classification’ trials, and (iv) ‘different-hand sound-pitch-classification’ trials. (D-K) Regression and psychometric curve results, shown separately for the four consecutive-trial types (i-iv). D,F,H,J, The coefficients of the previous stimulus (*s*_*t*−1_) and choice (*c*_*t*−1_) factor and their interaction terms with the granularity factor (*s*_*t*−1_ × *G*_*t*−1_; *c*_*t*−1_ × *G*_*t*−1_) are shown for the four consecutive-trial types. The filled circles indicate the coefficients that significantly deviate from 0 after control for FDR. E,G,I,K, The psychometric curves, shown for the four consecutive-trial types. The proportion of ‘greater’ choices, which belong to the upper half of the classes, was conditioned on *s*_*t*_ and *c*_*t*−1_ (upper) or *s*_*t*−1_ and *c*_*t*−1_ (lower) for each granularity condition. The color hues of the symbols and lines indicate the class of the previous choice. The circles and lines represent human choices and the predictions of the regression model, respectively.

In the last experiment (Exp 7), we tested this prediction by dissociating the DV and action spaces. Specifically, the action space was divided into two subspaces as we did for Exp 3: left-hand vs. right-hand spaces. One of the two subspaces was randomly used on a trial-to-trial basis by instructing observers which subspace to act upon with a pre-cue (Figure 7C). Unlike Exp 3, Exp 7 fixed the stimulus modality within a block and alternated it between blocks. If the granularity effect occurs in the action space, the anti-granularity effect is expected when hands are different between trials because the ‘large’ choice action with left-hand (‘3’ or ‘4’ response keys) would be closer to the ‘small’ choice action with right-hand (‘5’ or ‘6’ response keys) as granularity increases (Figure 7B).

The results support the DV-space account. We found that granularity modulated the choice-attraction bias, regardless of whether the action space was the same 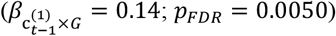 or different 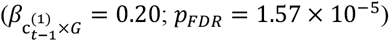 between two consecutive trials on the size task (Figure 7D-G). In the pitch task, it was less clear whether the granularity effect originates from the DV or the action space, as the granularity effect was insignificant when the action space differed 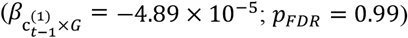, even though the granularity effect was significant when the action space remained consistent 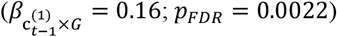 (Figure 7H-K). Based on these results, we conclude that the granularity effect cannot be ascribed to either action space shaping or commitment to the action space, and that the granularity effect based on the DV space is particularly robust in the size task.

### A Bayesian account of the history effects in reproduction, scaling, and classification tasks

Led by the idea that human decision-makers not just form abstract beliefs in the DV space but also granularize and update them over sequential episodes, we have conducted seven experiments demonstrating its implications. Next, we aim to provide a principled and unified account of the array of history effects observed in those experiments by translating this idea into a mathematical algorithm using Bayesian formalism^65-68^. These history effects can be summarized comprehensively by the 12 regression coefficients that capture how the four history factors in the previous classification-task trial—(i) choice (*c*_*t*−1_), (ii) modulatory granularity-on-choice (*c*_*t*−1_ × *G*_*t*−1_), (iii) stimulus (*s*_*t*−1_), and (iv) modulatory granularity-on-stimulus (*s*_*t*−1_ × *G*_*t*−1_) factors—influence the current responses in the three tasks—(i) reproduction, (ii) scaling, and (iii) classification tasks (Figure 8A). Thus, a valid model should be capable of simultaneously capturing all these 12 observed coefficients with a single set of parameters.

**Figure 8.**
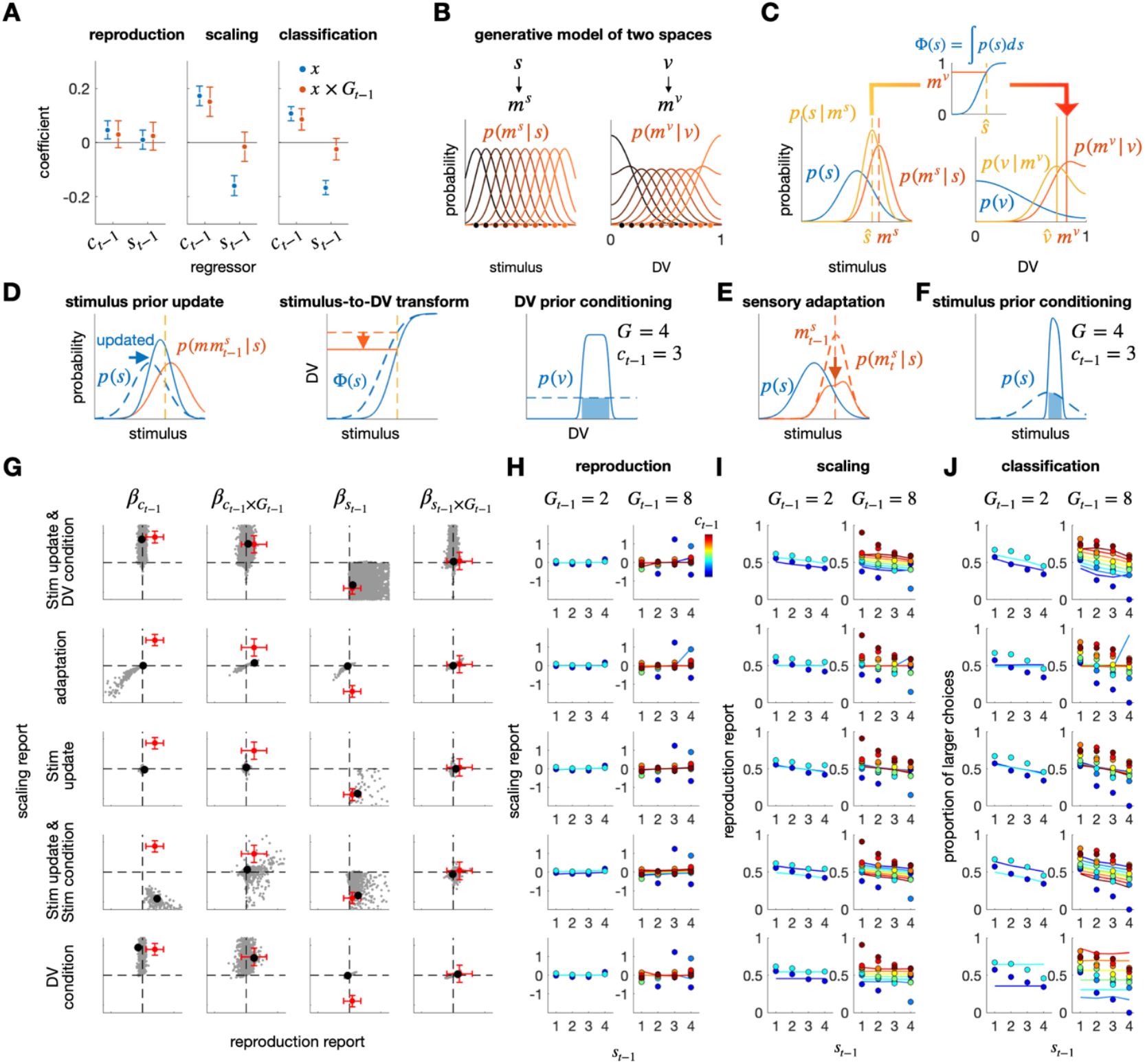
Results of the model simulations. (A) Benchmark history effects to be explained. Seeking a principled and unified account of the history effects found in the current study, we focused on the impacts of the previous choice (*c*_*t*−1_) and stimulus (*s*_*t*−1_), as well as their interaction with the previous granularity level (*c*_*t*−1_ × *G*_*t*−1_ and *s*_*t*−1_ × *G*_*t*−1_) on the current response for each of the three tasks: reproduction, scaling, and classification. Specifically, the regression coefficients found in the combined data from Exp 5 and 6 were used for the reproduction and scaling tasks, while those from Exp 4 were used for the classification task. Any successful account should simulate these 16 regression coefficients. The error bars correspond to the 95% bootstrap confidence intervals. (B) Internal models for stimulus and DV spaces. In the stimulus space, given a specific state of the stimulus variable *s*, a sensory measurement of the stimulus *m*^*s*^ is stochastically generated to form a likelihood function centered around the true state of *s* (left). In the DV space, given a specific state of the DV *s*, an internal measurement of the DV *m*^*v*^ is stochastically generated to form a bounded ([0, 1]) likelihood function centered around the true state of *v* (right). (C) Inferences in stimulus and DV spaces. The state of the stimulus variable is inferred given *m*^*v*^ as ŝ according to Bayes rule with the internal model described in panel B (left). The stimulus estimate in the stimulus space ŝ is transformed into a DV measurement *m*^*v*^, which is defined as the quantile of ŝ given the prior belief about the stimulus *p*(*s*) (middle). Given *m*^*v*^, the state of the DV is inferred as *v*4 according to Bayes rule with the internal model described in panel B (right). (D) Belief updating in the standard model (see main text). The prior beliefs are updated in parallel in both stimulus and DV spaces. The stimulus prior is updated to be attracted (from dashed to solid blue curves) toward the memory of previous stimulus (red curve) (left panel). As a result, the same stimulus estimate (vertical dashed line) is transformed into different states of *m*^*v*^ before and after the updating (from dashed to solid horizontal curves and lines in the middle panel). In the DV space, the DV prior is conditioned by the previous decision commitment, with varying extents based on its granularity (from dashed to solid curves in the right panel). (E) Changes in likelihood function in the *sensory adaptation model*. Sensory adaptation to the previous stimulus *s*_*t*−1_ leads to the distortion of the likelihood function in the current trial, which is suppressed (from dashed to solid red curves) around the previous stimulus measurement *m*^*s*^ (dashed vertical red line). (F) Stimulus prior updating in the *stimulus-prior-updating-by-the-choice model*. Unlike the standard model, this model assumes that the stimulus prior is updated to be conditioned by the previous decision commitment, with varying extents based on its granularity (from dashed to solid curves). (G-J) Simulation results from the standard model (top row) and its four variants (bottom four rows). G, Coefficients for the regression of the reproduction and scaling responses onto the stimulus (*s*_*t*−1_) and choice (*c*_*t*−1_) factor and their interaction terms involving the granularity factor (*s*_*t*−1_ × *G*_*t*−1_; *c*_*t*−1_ × *G*_*t*−1_) in the previous classification-task trial. In each panel, the coefficients for the scaling-task trials are plotted against those for the reproduction-task trials: gray dots, coefficients from individual simulations with different sets of model parameters (see STAR methods for details); black dots, coefficients from the simulations with the model parameters giving the outcomes closest to the observed ones, which are indicated by the red dots. H-J, Comparison of the best simulation outcomes to the observed ones. The reproduction (H) and scaling (I) responses of Exp 5 and 6, and the classification response (J) of the *G*_*t*_ fixed condition of Exp 4 are juxtaposed with the responses of models simulated with the parameters that best matches to the observed regression coefficients (black dots in G). The circles and lines represent responses of humans and models, respectively. The format of psychometric curves is consistent with the previous figures.

As a principled approach, we adopted the framework of Bayesian network modeling, which allows for belief updating across trials. Our central assumption is that decision-makers must form and update beliefs separately and in parallel within the stimulus and DV spaces to perform the three tasks normatively. This is because the variable to be inferred in the reproduction task differs from that in the other two tasks. One must infer the state of a physical variable (e.g., the physical size or location of a ball to catch or the physical melody of a song to sing) in the reproduction task, while one must infer the state of an abstract variable (e.g., the relative size or location of a ball compared to other balls in a reference group or the relative pitch of a note compared to other notes within a given melody structure) in the classification and scaling tasks. Another critical assumption is that the stimulus and DV spaces are separate yet “linked,” as decision-makers must convert values within the stimulus space—inferred states of a physical variable in the reproduction task—into the corresponding values within the DV space. This conversion is essential for optimally performing the scaling and classification tasks, which require calculating the probability of being greater than any other items in the distribution. Due to this hierarchical feedforward relationship from the stimulus space to the DV space, the reproduction task involves probabilistic inference only in the stimulus space, while the scaling and classification tasks involve probabilistic inference in both spaces. Critically, however, the stimulus prior—the belief formed and updated in the stimulus space—is used in that conversion but vanishes in the DV space because this conversion nullifies (flattens) any forms of priors in the stimulus space, which is a hallmark of the probability integral transformation in statistics. This nullification of the stimulus prior during the conversion necessitates the DV prior in the DV space.

We implemented these assumptions in our Bayesian network model by positing that the stimulus and DV spaces have their own generative models where a stimulus measurement (*m*^*s*^) is stochastically generated from the stimulus variable (*s*) in the stimulus space while a DV measurement (*m*^*v*^) is stochastically generated from the DV (*v*) in the DV space (Figure 8B). Consequently, two independent prior beliefs can propagate across trials in parallel within the stimulus and DV spaces. This parallelism in belief propagation between the two separate representational spaces is key to explaining why and how the stimulus-repulsion and the choice-attraction biases occur in both scaling and classification tasks but not in the reproduction task, as elaborated below.

From these two generative models, normative inference processes are derived to perform the three tasks (Figure 8C). The inferential processes have three modules: the *s*-space module, where beliefs about stimulus states (e.g., absolute magnitudes of size) are formed and updated (the left panel of Figure 8C); the *v*-space module, where beliefs about DV states (e.g., probability of being larger than others in size) are formed and updated (the right panel of Figure 8C); and the *s*-to-*v* transformation module, which nonlinearly maps beliefs about the stimulus states to the DV states using the cumulative distribution function of the stimulus prior (Φ) (the center panel of Figure 8C). With these modules, the three tasks are carried out as follows. The reproduction task is carried out within the *s*-space module because it requires inferring the absolute magnitude of a stimulus, which is obtained by calculating the maximum a posteriori (MAP) of stimulus 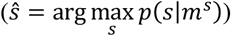 based on the stimulus prior *p*(*s*) and likelihood function *p*(*m*^*s*^|*s*) (the left panel of Figure 8C). Then, *m*^*v*^ (the DV measurement) results from the *s*-to-*v* transformation, via which the inferred state of the stimulus ŝ is mapped onto the DV space. The scaling and classification tasks are both carried out in the *v*-space module because they require inferring the relative magnitude of a stimulus, the DV. For the scaling task, a point estimate of DV is obtained by calculating the MAP of DV 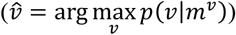, which involves combining the DV prior *p*(*v*) and likelihood function *p*(*m*^*v*^|*v*) in the *v*-space module (the right panel of Figure 8C). For the classification task, the DV space is evenly discretized according to the granularity, and a class is chosen as the order of the discretized range where 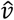 is located.

In what follows, we provide an intuitive account of how our model can predict the history effects observed in the three tasks. As our group previously explained^46,49,50^, when a stimulus is observed, the stimulus prior *p*(*s*) is updated to be attracted to the memory of that stimulus (*mm*^*s*^) (left panel of Figure 8D). Because of this update in *p*(*s*), the transformation function is also attracted toward the previous stimulus. In turn, this attracted transformation function causes the DV in the subsequent trial to be repelled away from the previous stimulus (center panel of Figure 8D). Thus, the larger the previous stimulus, the smaller the current DV measurement, and the more the current response is repulsed away from the previous stimulus. This explains the stimulus-repulsion bias 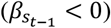 observed in both classification and scaling tasks (the two right panels of Figure 8A), while it does not induce the stimulus-repulsion bias in the reproduction task because what is repelled is not the stimulus itself but a DV measurement (left panel of Figure 8A). Furthermore, it does not induce any granularity effect on the stimulus-repulsion bias because the granularity was not incorporated into the stimulus prior updating.

As for the choice-attraction bias and granularity effect, we posit that when a decision is made, the posterior belief of DV is narrowed down to match the range of DV that the decision commits to. This belief is then carried over into the future, serving as the DV prior in the next trial (right panel of Figure 8D). Thus, the more granular the previous decision, the narrower the current DV prior, and the more the current response is attracted to the previous decision. This explains the choice-attraction bias 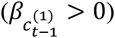 and the granularity effect 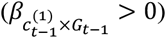 in both classification and scaling tasks (the right two panels of Figure 8A). In the reproduction task, neither the choice-attraction bias nor the granularity effect occurs (the left panel of Figure 8A), as what is granularized is the DV space, while an estimate of size magnitude does not require accessing the DV space.

To confirm our intuitive account, we conducted model simulations^50,69^ using a range of model parameters that were deemed reasonable based on participants’ performance in Exp 4-6 (see STAR methods for details). To assess the history effects in the model’s choice behavior, we performed the multiple regression analysis and summarized the results using conditioned psychometric curves, following the same procedure as with human participants. The simulated regression coefficients show that our model’s behavior (the gray dots in the top panel of Figure 8G) can display the pattern of history effects that are similar to those observed for human participants (the red dots in the top panel of Figure 8G). When the best model parameters were selected, the pattern of historical effects showed remarkable similarity between the model and human data (as the colored lines in the top panels of Figure 8H-J).

### Falsification of alternative accounts of the observed history effects

Having confirmed our Bayesian network model’s capability to produce the observed history effects, we now investigate whether these effects can be produced by alternative scenarios that do not necessarily entail creating the DV space and updating beliefs across episodes there, which is central to our explanation of the observed history effects.

First, we considered the ‘sensory-adaptation’ hypothesis as an alternative scenario to the ‘stimulus-prior-updating’ of our model in explaining the stimulus-repulsion bias. This hypothesis posits that the previous stimulus reduces the sensory apparatus’s encoding gain, potentially causing the current perception to be repelled away against the previous stimulus^43,70,71^. We incorporated the ‘sensory-adaptation’ scenario into our model by modifying the stimulus likelihood function^72^ (as shown in Figure 8E; see STAR methods for details). This scenario can generate the stimulus-repulsion bias in the scaling task 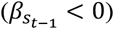. However, whenever it does, it also generates the same bias in the reproduction task (the second row of Figure 8G-J), which is an uncorrectable deviation from the observed effects.

Second, the failure of ‘sensory adaptation’ to capture the stimulus-repulsion bias in the scaling task suggests that it may arise from a different source. Thus, we considered the ‘stimulus-prior-updating-by-the-stimulus’ hypothesis (see STAR methods for details). This scenario can generate all history effects of the previous stimulus in the three tasks simultaneously (the symbols involving *s*_*t*−1_ on the third rows of Figure 8G-J). Particularly, it can simulate the stimulus-repulsion bias during the scaling and classification tasks without displaying it in the reproduction task. However, it cannot display any history effects of the previous choice (the symbols involving *c*_*t*−1_ in the third rows of Figure 8G-J), and therefore cannot be considered a comprehensive explanation for the observed history effects.

Third, despite its failure to capture the previous choice effect, the effectiveness of the stimulus-prior-updating-by-the-stimulus hypothesis in explaining the previous stimulus effect suggests that stimulus-prior-updating may account for the history effects if it is appropriately modified to incorporate the previous choice effect. Thus, we considered the ‘stimulus-prior-updating-by-the-choice’ hypothesis as a scenario that granularizes and updates beliefs over episodes, not in the DV space, but in the stimulus space^73,74^ (Figure 8F; see STAR methods for details). Since it is adapted from the stimulus-prior-updating-by-the-stimulus model, the stimulus-prior-updating-by-the-choice hypothesis can generate the history effects of the previous stimulus. (the symbols involving *s*_*t*−1_ on the fourth rows of Figure 8G-J). However, it still cannot produce the effects of the previous choice: it displays the choice-repulsion bias, instead of the choice-attraction bias, and fails to generate the granularity effect (the symbols involving *c*_*t*−1_ on the fourth rows of Figure 8G-J). Thus, the ‘stimulus-prior-updating-by-the-choice’ hypothesis does not comprehensively account for the observed history effects.

Finally, one might speculate that the skewed likelihood distribution in the DV could induce the repulsive bias without updating the stimulus prior, as a previous study demonstrated that skewed likelihood distribution induced the repulsive bias from the stimulus prior^75^. Thus, we examined whether the skewed DV likelihood and the belief updating in the DV space can reproduce both the granularity effect and the stimulus-repulsion bias without updating the stimulus prior (Methods). However, the ‘DV-prior-updating-by-the-choice’ could not reproduce the stimulus-repulsion bias, even though it can simulate the choice-attraction bias and its modulation by decision granularity (on the fifth row of Figure 8G-J). Therefore, the repulsive bias could not be reproduced solely by the skewed likelihood of the DV-space updating.

As we have shown earlier, when we modified the ‘stimulus-prior-updating-by-the-stimulus’ model by hypothesizing that belief is granularized and updated in the DV space rather than in the stimulus space, all of the history effects were successfully captured. This success, along with the failure of the alternative hypotheses, to generate and produce the observed history effects, underscores the necessity for dual belief propagations through both the stimulus space and the DV space.

## DISCUSSION

The essence of human cognition, particularly its most advanced form in comparison to other natural and artificial intelligences, lies in the ability to give abstract structure to daily experiences^76,77^. A prime example of structured abstraction in human cognition is forming DVs from limited or impoverished sources, including sensory evidence, prior knowledge, and action cost (reward) to guide decision-making with uncertainty^2,9,14^. The processes involved in DV formation have been studied extensively, but the utilization of DVs after formation remains relatively unexplored despite its importance in adapting to the environment. Based on the concept that the space representing DVs is well-suited for efficiently translating past experiences into future expectations, we hypothesized that (i) when a categorical decision is made, it restricts the belief distribution about DV states to a particular range in the DV space, and (ii) this restricted belief distribution is then carried forward to future trials and used as a prior expectation for potential DV states. As an empirically testable implication of this hypothesis, we predicted the granularity effect: the choice-attraction bias—the extent to which the current choice is attracted toward the previous one—increases as the commitment to DV states becomes finer with an increasing level of decision granularity.

Confirming this prediction, human individuals reliably displayed the granularity effect. Critically, its presence and absence in our experiments suggest that it originates from the DV space. To summarize, the granularity effect: (i) *did* occur when size (pitch) classification or scaling was followed by size (pitch) classification regardless of whether different hands were used for enacting the previous and current choices; (ii) *did not* occur when size classification was followed by pitch classification, and vice versa; (iii) *did not* occur when size classification was followed by size reproduction; (iv) *did not* occur on the stimulus-repulsion bias. These results suggest that the granularity effect does not originate from the action space (i), the source-general DV space (ii), nor the stimulus space (iii and iv), but rather from the source-specific DV space (black and gray arrows with green shades in Figure 9).

**Figure 9.**
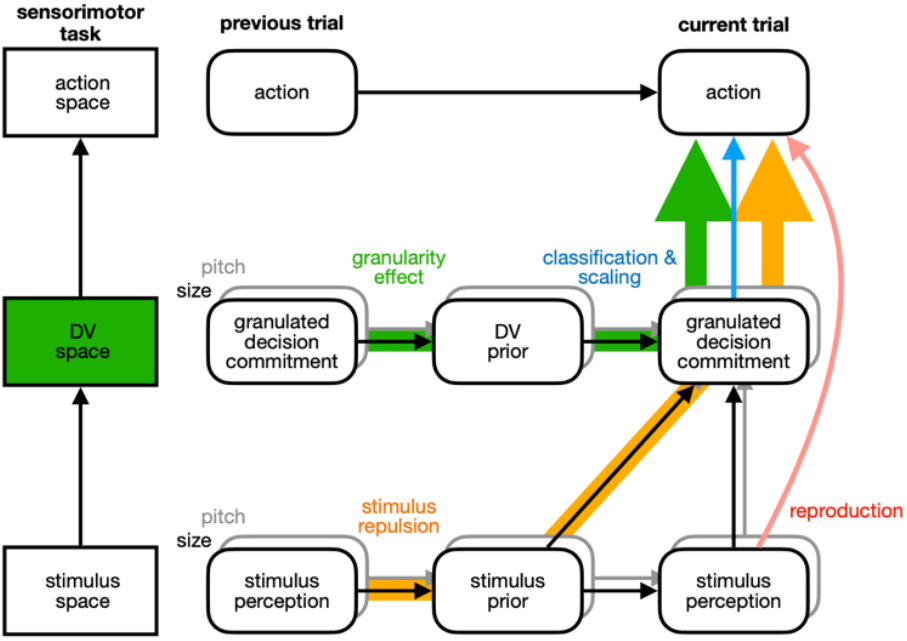
Distinct routes of belief propagation for the history effects in the reproduction, scaling, and classification tasks. The cognitive acts involved in the three tasks are summarized in three layers of spaces: stimulus perception in stimulus space (bottom), decision commitment in DV space (middle), and action in action space (top). Between consecutive trials, prior beliefs are updated in parallel in both stimulus and DV space. According to our proposed standard model, the stimulus-repulsion bias originates from the route starting with stimulus perception in the previous trial, then the update of the stimulus prior, and finally the decision commitment in the current trial (arrows with yellow shades). The choice-attraction bias and its modulation by granularity (the granularity effect) originates from the route starting with the decision commitment in the previous trial, then the update of the DV prior, and finally the decision commitment in the current trial (arrows with green shades). As a result, all these history effects do occur in the scaling and classification tasks, which require accessing the DV space (blue arrow), but the granularity effect and choice attraction do not occur in the reproduction task, which does not require accessing the DV space (pink arrow).

As a principled and unified account of the history effects observed under the various task conditions in our experiments, we built a Bayesian model in which probabilistic beliefs propagate over episodes in both stimulus and DV space. The model’s ability to produce the observed history effects suggests that the human mind does form beliefs with varying degrees of uncertainty not just in the stimulus space but also in the DV space. Furthermore, it suggests that these beliefs are used for future purposes through probabilistic computation in the stimulus and the DV spaces in parallel.

Our findings expand the current understanding of human cognition by showing that humans translate their experience into a *generalizable* form of knowledge during categorical decision-making, form its probabilistic representation in an *abstract* continuous space that is *separate* from where sensory stimuli or motor plans are represented, and update and apply this continuous belief distribution over episodes of cognitive acts with different sensorimotor specifics to adjust their behavior trial-to-trial in ever-changing environments effectively.

### Consequences of decision commitment

Previous studies on decision-making focused on how decision-makers form DVs and commit to a specific option based on their state. Signal detection theory^1^ (SDT) has been instrumental in guiding research on DV formation and decision rules by defining the DV as the ratio of likelihoods of binary options and offering a decision rule based on criterion setting. Sequential analysis theory^3,4^ (SAT) adds a dynamic quality to DV formation and offers a decision rule to stop DV formation and commit to a decision.

Guided by SDT and SAT, systems neuroscientists significantly advanced our understanding of decision-making by identifying neural correlates of DV state^18,78-81^, dynamic DV formation^82-84^, and decision commitment^16,85^, despite being challenged by recent perspectives and findings (see ^86,87^ for extensive reviews). We took a step further by exploring the cognitive consequences of decision commitment. By examining the impact of decision granularity on future cognitive processes, we have discovered that the belief about the DV state is updated as a consequence of decision commitment and is subsequently integrated into the future formation of DV as a form of prior belief. Specifically, the granularity effect suggests that committing to a choice restricts the belief distribution to a discrete range of the DV state linked to that choice.

*Why* does the brain restrict the belief of DV in accordance with granularity? We conjecture that this belief restriction has a rational basis. Making a decision is a deliberate process that requires significant mental effort^88,89^ and involves a hierarchy of brain regions^87,90-93^, which uses up substantial neural computation and communication resources^94,95^. In addition, the quality of decisions would likely improve with an increasing amount of resources^35,96,97^. This means that as a decision becomes more granular, more resources are spent to commit (as evidenced in longer RTs; mean RTs of Exp 1: 0.65 s, 0.82 s, and 0.96 s for *G* = 2, 4, and 8 respectively; Exp 2: 0.74 s, 0.89 s, and 1.04 s for *G* = 2, 4, and 8 respectively). Therefore, giving more weight to decisions with finer granularity seems reasonable, reinforcing beliefs for which more resources are spent. In this sense, the granularity-dependent belief restriction of DV can be viewed as an instantiation of resource-rational computation^96,98-100^.

### Shaping beliefs within the decision-variable space

In the context of probabilistic knowledge^101-104^, “belief” refers to a subjective knowledge of probabilistic information. Beliefs are adaptive, and therefore normative, as long as they accurately reflect the actual states of variables and their true relationships within the environment. Therefore, a key aspect of explaining adaptive behavior is understanding how individuals form and use their beliefs to adapt to their surroundings.

Probabilistic decision theories have been offering powerful accounts of how human decision-makers carry out various sensory-motor tasks by positing that they can form nearly optimal beliefs about the state of the world (prior knowledge)^105-109^, the contingency between sensory evidence and its causal state of the world (likelihood function)^75,110^, and the contingency between the motor command and its outcome in the world (conditional distribution)^73,74^, and the reward linked to a particular outcome (loss function)^110^. It should be stressed that these beliefs are all assumed to be formed about the states and outcomes of the concrete world, i.e., stimulus measurements or motor commands (e.g., ‘motion direction and its sensory signals’ or ‘pointing plan and its hand location in world coordinates’). However, DVs have never been treated as a random variable with uncertainty but only as a point estimate without uncertainty^1,9,111,112^. Our simulation of different versions of the Bayesian model revealed that to display the granularity effect, the probabilistic belief in the DV space needed to be formed and updated through decision commitment, resulting in the restriction of the belief to a specific range of the DV state corresponding to the *granulated* commitment. These results suggest that beliefs are updated in parallel, with one in the stimulus space engendering the stimulus-repulsion bias (arrows with orange shades in Figure 9), and the other in the DV space engendering the choice-attraction bias and granularity effect (arrows with green shades in Figure 9).

Our interpretation above raises a normative question: *Why* does the human brain shape probabilistic knowledge in the DV space rather than only in the stimulus or action spaces? In other words, what benefits does updating beliefs in the DV space offer? We conjecture that several points can justify such belief shaping. First, beliefs in the DV space can be more invariant to nuisances and thus be more generalizable compared to those in the stimulus or action space. For instance, the absolute size and brightness of a thing are subject to changes depending on the viewing distance and illumination, respectively. However, its size and brightness relative to those of the others in a population are tolerant to such nuisance factors. Likewise, enacting a decision in a specific part of space is subject to changes depending on the provisional contingency between choice and action policy. In such circumstances, it would be more reasonable to hold beliefs in a representational space immune to such contingencies. This makes the DV space a reliable place to store knowledge from past decision-making episodes and use it for future episodes. Second, the prior belief about stimulus distribution becomes, in principle, annulled in the DV space. This is because the DV measurements are linked to the stimulus estimate via the cumulative distribution functions of the stimulus prior belief (Figure 8C). Suppose the stimulus estimates ŝ follow a distribution similar to the stimulus prior *p*(*s*). Then, the transformed DV measurements *m*^*v*^ = Φ(ŝ) would be uniformly distributed in the DV space. Hence, the DV space needs its own prior beliefs to utilize the probabilistic information that relates past to future events in an abstract manner. Lastly, DVs contain rich, task-relevant information in a concise format because they are formed by combining multiple factors that help achieve the defined objective of the task. This makes the DV space an efficient place to form and propagate beliefs over a sequence of cognitive events. In summary, the DV space offers a general platform for propagating probabilistic knowledge over a sequence of decision-making episodes, even in complex environments filled with distracting factors in the stimulus and action domains.

Our Bayesian account of the granularity effect based on probabilistic computation in the DV space leads to an empirically testable prediction about the neural representation of DVs. Specifically, our account suggests that the neural signals of DVs should contain information about the uncertainty regarding the trial-to-trial states of the task-relevant DVs. Recently, human neuroimaging^113^ and monkey electrophysiological^114^ studies have shown that population neural activities in the primary visual cortex carry uncertainty-associated signals in the stimulus representation space (e.g., visual orientation feature). Thus, one may take a similar approach to the high-tier cortical regions where DVs are likely to be represented, such as the parietal or prefrontal cortices. We predict that their trial-to-trial population activity conveys uncertainty representations about DVs that better explain decisions than point estimate representations. Notably, our Bayesian account of the granularity effect suggests that the level of uncertainty carried by the DV-associated neural signals decreases as decision granularity increases.

### Distinguishing between the perceptual history effects arising in the stimulus and DV spaces

The phenomenon of history effect, also known as “serial dependence,” has been observed in various perceptual tasks (see ^115^ for a comprehensive review). The most commonly reported type of history effect is the positive sequential dependence, also known as “attractive bias,” where various cognitive processes, including estimation and decision-making, are biased toward recent perceptual experiences^116^. The attractive bias has been observed in two tasks: reproduction and categorization tasks. In the former, observers need to reproduce a target’s feature on a continuous feature spectrum using an adjustment method^44,47,55,117-120^, as was done in the ‘reproduction-task’ trials of Exp 5 and 6 in the current work. In the latter, observers decide which particular ordinal category a target’s feature value belongs to using a forced-choice method^56-59,121^, as was done in the ‘classification-task’ trials of all the experiments in the current work.

It is unclear whether “attraction” arises from the same source when performing reproduction and categorization tasks^56,115,116^. In a recent review^115^, the differences in history effect between the two tasks were attributed to the difference between continuum or discrete values used for reporting. However, our findings indicate that the difference in representational space is the critical distinction between reproduction and categorization tasks, which require accessing the stimulus and DV spaces, respectively. Supporting this idea, we found that the choice-attraction bias and the granularity effect were present when the classification-task trial was followed by the scaling-task trial but not when followed by the reproduction-task trial (Exp 5 and 6). This points out that the key factor in transferring past experiences into future ones is not the consistency between consecutive trials in reporting format (whether discrete or continuous), but rather the commonality of their representational space.

The other history effect observed in the current work is the stimulus-repulsion bias. Critically, similar to the choice-attraction bias, the stimulus-repulsion bias occurred when the classification trial was followed by the scaling trial but not when followed by the reproduction trial. These results help to resolve the issue about the origin of the stimulus-repulsion bias in the classification task, determining if it arises from sensory adaptation^43,70,115,122^ or from updating stimulus distribution^45,46,49,50,54^. The sensory adaptation hypothesis implies that the stimulus-repulsion bias occurs regardless of whether the classification trial was followed by the scaling trial or by the reproduction trial because the key factor for sensory adaptation is the physical stimulus itself. Our findings defy this implication and suggest that updating the stimulus distribution is the source of the stimulus-repulsion bias in the classification task. This conclusion corroborates the conclusion drawn from a recent neuroimaging study of our group^46^.

### Potential confounds of decision granularity

We found that the granularity of the previous decision *modulates* the extent to which the current choice is biased toward the previous one. We attributed this modulatory effect of decision granularity to belief updating in the DV space. Could this modulation be caused by other modulatory factors that are known to influence serial dependence, such as ‘stimulus uncertainty,’ ‘performance confidence,’ and ‘attention’? We are opposed to this possibility for the following reasons.

Previous studies demonstrated the modulatory influence of stimulus uncertainty on serial dependence by manipulating the orientation cardinality^41,123^, spatial frequency^41,124^, contrast^125^, eccentricity^70^, and noise^126^ of visual stimuli. In our study, these stimulus parameters remained constant and, therefore, did not covary at all with decision granularity. Hence, the granularity effect is unlikely to originate from stimulus uncertainty.

While the physical properties of stimulus input objectively determine stimulus uncertainty, performance confidence subjectively varies from trial to trial^36,127^. Previous studies have shown that a higher level of performance confidence in the previous trial tends to promote serial dependence with a stronger attractive bias^128,129^ and that the proxy measures of confidence, like RT and pupil size, also influence the degree of serial dependence^56-59^. However, the modulatory direction of this previous confidence is opposite to the granularity effect because, although higher levels of confidence are supposed to promote the choice-attraction bias^56,59^, the RT data indicate that the level of confidence becomes lower as the level of decision granularity increases. Furthermore, even when we included the RT measurements as the modulatory factors for the influences of the previous stimuli and choices when evaluating the granularity effect using the multiple regression model, the granularity effect could not be explained away by changes in RT from trial to trial.

In previous studies, researchers manipulated the spatial^130^ and featural^131^ aspects of attention, as well as the amount of attention, by instructing participants to either report or not report stimulus features^70,132^. By doing so, they showed that the attractive serial dependence increases with proper attention to the previous stimulus. This raises the question of whether the level of decision granularity was confounded with attention. However, this ‘attention’ scenario cannot explain why the granularity effect only occurred when the classification-task trial was followed by the scaling-task trial, and not when it was followed by the reproduction-task trial (Exp 5 and 6). If the effect were caused by attention, it should have been observed in both conditions.

Besides these modulatory factors of history effects, one might consider that the granularity effect may share its origin with a phenomenon known as ‘choice hysteresis’^133-135^. According to this view, the granularity effect is not mediated by belief updating in the DV space but arises from the system’s inability to make a quick state transition between trials^133-135^, resulting in a failure to disengage from the previous decision-making episode. However, this interpretation is unlikely for the following two reasons. First, the phenomenon of choice hysteresis typically occurs when a variable of interest gradually shifts its state, indicating that its current state is closely tied to the previous state. However, in our study, the current state of the variable of interest (i.e., the class of visual size or sound pitch) was completely dissociated from its previous state, as its state was randomized from the previous one. Second, the phenomenon of choice hysteresis is known to decrease as the inter-trial interval (ITI) increases. Therefore, if the granularity effect shares its origin with choice hysteresis, it should also lessen as a function of ITI. We examined this implication using one of our group’s datasets obtained from a different experiment, where both ITI and decision granularity were varied on a trial-to-trial basis (Figure S1A). This experiment confirmed the granularity effect, demonstrating its robustness. However, the impact of ITI differed from the findings related to choice hysteresis: the granularity effect was more pronounced in trials with longer ITI compared to those with shorter ITI (Figure S1B,C). Considering the independence of the stimulus sequence from trial to trial and the influence of ITI on the granularity effect, we conclude that the granularity effect does not share its origin with choice hysteresis.

### The granularity effect as a novel history effect

Given the robust yet highly specific nature of the granularity effect, we conducted a thorough search of the existing literature to identify any previous mentions of comparable phenomena. To our best knowledge, a kind of sequential effect most similar to the granularity effect has been observed in a series of studies focusing on the ‘absolute identification’ task^136-139^. In this task, where participants needed to report the exact magnitude of a stimulus feature, the influence of the previous report on the current one tended to grow as the number of response options increased. However, this increase in sequential dependence cannot be solely attributed to the number of response options, as the number of stimuli was perfectly correlated with that of response options in the absolute identification task. When they were decorrelated in one exceptional study^137^, there was no significant increase in sequential dependence as a function of response option. Newly registering the granularity effect in the list of history effects, we assert that any comprehensive account of the history effects must address the granularity effect.

## LIMITATION OF STUDY

As a computational-level account^140^ of the granularity effect, we proposed a Bayesian model. In this model, beliefs are formed and updated according to a generative model that assumes representational stability in the DV space. Although this model successfully captured all the history effects observed under various experimental conditions, it has limitations that need to be addressed to provide a comprehensive account of the history effects. For sure, the model needs to be extended to incorporate other decision-making tasks that may require DV spaces different from the one needed to classify stimulus ordinality, as in our experiments. Such an extension would require probing the granularity effect in alternative decision-making tasks involving a varying number of choice options, particularly those that do not depend on the order of stimuli^141-143^.

Despite this need for further elaboration, our findings point to the DV space as a previously unrecognized source of trial-to-trial belief updating. We believe that future efforts to account for the across-trial belief updating processes in the DV space will significantly advance the current understanding of how the human brain efficiently adapts to its surroundings by constructing an abstract and continuous —thus highly generalizable—belief distribution out of their categorical decisions.

## ACKNOWLEDGEMENTS

This research was supported by the National Research Foundation of Korea funded by the Ministry of Science and ICT (RS-2024-00435727, RS-2024-00349515, RS-2023-00276729).

## AUTHOR CONTRIBUTIONS

Conceptualization, H.L., J.L. and S.-H.L.; Methodology, H.L. and S.-H.L.; Investigation, H.L. and S.-H.L.; Writing – Original Draft, H.L.; Writing – Review & Editing, H.L., J.L. and S.-H.L.; Funding Acquisition, H.L. and S.-H.L.; Resources, S.-H.L.; Supervision, S.-H.L.

## INCLUSION AND DIVERSITY

We support inclusive, diverse, and equitable conduct of research.

## Declaration of Generative AI and AI-assisted technologies in the writing process

During the initial drafting process, H.L. utilized ChatGTP to verify the naturalness of his phrases and sentences. Afterward, the corresponding author, S.-H.L., thoroughly revised the initial draft and made the necessary edits to the content. S.-H.L. takes full responsibility for the publication’s content.

## SUPPLEMENTAL INFORMATION

## RESOURCE AVAILABILITY

## Lead contact

Further information and requests for resources should be directed to and will be fulfilled by the lead contact Sang-Hun Lee.

## Materials availability

Besides data and MATLAB codes, this study did not generate any new reagents or materials.

## Data and code availability

- All data have been deposited at GitHub and are publicly available. The DOI is listed in the key resources table.
- All original code has been deposited at GitHub and is publicly available. The DOI is listed in the key resource table.
- Any additional information required to reanalyze the data reported in this paper is available from the lead contact upon request.

## EXPERIMENTAL MODEL AND STUDY PARTICIPANT DETAILS

The study consisted of eight experiments (Exps 1-7 and a supplementary experiment (Exp S)) with 100 human participants having normal or corrected-to-normal vision and hearing. The participants took part in one or more of these experiments, as follows: 58 participants (39 females; aged 18-33 years) in Exps 1 and 2; 18 participants (11 females, aged 22-33 years) in Exp 3; 18 participants (11 females, aged 20-33 years) in Exp 4; 11 (5 females, aged 19-31 years) participants in Exp 5; 15 participants (13 females, aged 19-29 years) in Exp 6; 30 (19 females, aged 20-33 years) participants in Exp 7; 16 (7 females, aged 19-26 years) in Exp S. Prior to participating, they all provided informed consent and were unaware of the purpose of the experiments. The experimental procedures were approved by the Research Ethics Committee of Seoul National University.

## METHOD DETAILS

### Main experimental setup

Visual and auditory stimuli were displayed on a computer monitor (LG 27UL550, refresh rate of 60 Hz and resolution of 1920 × 1080 pixels) and heard through a two-channel speaker (Altec Lansing Series 100) in a dimly lit enclosed space. The viewing distance was 62 cm and enforced with a chin rest. The codes for controlling experiments, including stimulus generation and presentation, were written in Matlab (Mathworks, Inc.) in conjunction with the Psychtoolbox^144^ and run on an Apple Mac Pro computer with Quad-Core Intel Xeon E5 3.7 GHz, 12 GM RAM. Manual responses were received by an i-rocks KR-6170 keyboard.

### Experiment 1

Exp 1 consisted of three types of blocks: ‘threshold-calibration,’ ‘training,’ and ‘testing’ blocks. In the threshold-calibration blocks, participants began each trial by fixating a small dot (diameter in visual angle, 0.27°; luminance, 0.98 cd/m^2^) appearing at the center of a dark screen (1.11 cd/m^2^), which remained throughout the trial to help them maintain their fixation. Half a second after the fixation dot appeared, a thin (standard deviation of Gaussian envelope, 0.11°), white (1.45 cd/m^2^) dashed (radial frequency, 30 cycles/360°) ring was presented for 0.3 s. Participants were required to classify the ring as ‘small’ or ‘large’ based on size within a 1.5-s decision window. We provided feedback for their choice on a trial-by-trial basis by changing the fixation dot’s color for 0.5 s to green, red, or yellow, depending on whether the choice was correct, incorrect, or a miss, respectively. The correctness was determined using the class boundary set at the 3° radius. The miss trial is a trial where participants failed to respond within the decision window or pressed an undesignated button.

For each participant, a tailored threshold difference in ring size was calibrated through 4 blocks of trials, each consisting of 50 trials, using three randomly interleaved chains of one-up-two-down staircases according to the PEST rule^89^, where the three chains started from the easiest, intermediate, and hardest stimuli, respectively. The first 20 trials were excluded from analysis for their instability. The cumulative normal distribution function was fit to the classification psychometric curve using a maximum-likelihood procedure. The threshold difference was defined as the ring size associated with a 90 percent correct proportion based on the fitted function. Ring sizes for the training and testing blocks of trials were sampled from a normal distribution with a mean corresponding to the pre-determined boundary size (3°) and a standard deviation determined by the threshold difference.

The threshold calibration blocks were initially followed by the alternating training and testing blocks for the first 18 blocks, and then solely by the testing blocks for the remaining 12 blocks. The structure of the training and testing blocks was the same as the threshold-calibration block, except that each block had 45 trials, and the number of choice options (classification granularity) ranged from 2, 4 to 8. For each block, the classification granularity was set at a single level randomly chosen from the three levels. Participants were informed of this before the block began. The training and testing blocks differed only in that trial-to-trial corrective feedback was provided only in the former. The feedback during the training blocks resembled the threshold-calibration blocks, but with the additional feature of displaying the correct response button in incorrect and miss trials. For each participant, a total of 315 trials were acquired for each granularity level across the 21 testing blocks (9 blocks from the alternating period and 12 blocks from the non-alternating period). The trials used for data analysis were exclusively from these testing blocks.

As mentioned earlier, the ring size on each trial of the training and testing blocks was chosen randomly from a distribution based on the results of the threshold-calibration trials. For any given ring size, its class was defined by the median, quartiles, and octiles of this size distribution for the granularity levels of 2, 4, and 8, respectively. It should be noted that the size distribution remained constant throughout the entire testing and training blocks, while the classification granularity varied from block to block. This allowed us to manipulate the main variable of interest, granularity level, while controlling for any potential effects of stimulus range.

Participants indicated their choice by pressing a number key with both hands on the computer keyboard (Figure 7A): the [4,5], [3,4,5,6], and [1,2,3,4,5,6,7,8] keys were used in the blocks with granularity levels of 2, 4, and 8, respectively. The participants were instructed to use a specific finger for pressing each key: the index, middle, ring, and little fingers were to be used for the [4,5], [3,6], [2,7], and [1,8] keys, respectively. The left hand was assigned to the [1,2,3,4] keys, and the right hand was assigned to the [5,6,7,8] keys.

### Experiment 2

In Exp 2, the procedure was identical to that of Exp 1, except that beep sounds were used instead of ring stimuli. The beeps were classified based on pitch rather than size, with a specific pitch boundary set at 170 Hz. The beep sounds were created using a Psychtoolbox function called ‘Beeper.m’. The volume of the sounds was adjusted individually to ensure that each participant could comfortably hear them throughout the entire experiment. The order of Exp 1 and 2 was randomized across participants: 26 and 32 of them performed Exp 1 and 2 first, respectively.

### Experiment 3

In Exp 3, eighteen of the individuals who participated in Exp 1 and 2 took part. On each trial, participants performed either the ring size classification task (the one used in Exp 1) or the sound pitch classification task (the one used in Exp 2). The task was randomly chosen for each trial and was indicated at the start of the trial by displaying the words “Ring” or “Pitch” on the screen. The level of classification granularity alternated between 2 and 4 on a block-to-block basis. For each participant, the ring sizes and sound pitches were sampled from the same tailored distributions used in Exp 1 and 2, respectively.

To prevent any factors associated with motor or action space from confounding the transfer of granularity effect between visual and auditory modalities, we made participants use the fingers of the different hands for the two tasks. In the blocks with the granularity level of 2, the [2,3] keys were pressed by the third and middle fingers, respectively, of the left hand for ring size classification while the [6,7] keys were pressed by the middle and third fingers, respectively, of the right hand for sound pitch classification (the row labeled by “*G* = 2” in Figure 7B). In the blocks with the granularity level of 4, the [1,2,3,4] keys were pressed by the ring, third, middle, and index fingers, respectively, of the left hand for ring size classification while the [5,6,7,8] keys were pressed by the index, middle, third, and ring fingers, respectively, of the right hand for sound pitch classification (the row labeled by “*G* = 4” in Figure 7B).

### Experiment 4

In Exp 4, eighteen of the individuals who participated in Exp 1 and 2 took part. The procedure of Exp 4 was similar to that of Exp 1. Participants performed the ring size classification task, beginning with alternating blocks of the training and testing trials, followed by a period of testing-trial-only blocks. Exp 4 aimed to determine the temporal locus of the granularity effect, specifically whether the choice-attraction bias increases as a function of the granularity in the previous trial or the current trial. To investigate this, Exp 4 included two types of trial blocks: one with fixed granularity at the level of 2 and the other with varying granularity between two levels, 2 and 8. Comparing the trials between the two types of blocks allowed us to create two critical conditions: one in which the granularity varied in the previous trial but fixed in the current trial (marked by the orange dashed box and letters in Figure 5), and the other in which the granularity fixed in the previous trial but varied in the current trial (marked by the green dashed box and letters in Figure 5). Exp 4 began with the alternating training and testing blocks for the first 16 blocks, which were followed by the testing blocks for the remaining 20 blocks, each block consisting of 50 trials.

To address potential confounds related to cognitive load in switching action maps, we carefully assigned keys to choices in the following way. In the blocks with fixed granularity, we alternated key assignments between the [c,v] keys and the [4,5] keys (as shown in the bottom panels of Figure 5). In the blocks with varying granularity, we alternated key assignments between using the [c,v] keys for the granularity level of 2 and the [1,2,3,4,5,6,7,8] keys for the granularity level of 8 (as shown in the top panels of Figure 5). The fingers were assigned to the keys in the same way as in Exp 1, except that the left and right thumbs were assigned to the [c,v] keys.

### Experiment 5

Exp 5 aims to determine if the granularity effect can be generalized across different perceptual tasks depending on whether the tasks require the same decision variable (DV). Three different tasks were performed on ring stimuli: (i) classification task, (ii) scaling task, and (iii) reproduction task. The overall procedure of the classification task was similar to that used in Exp 1, 3, and 4. In the scaling task, participants were shown a ring and asked to judge which quantile its size belongs to within the entire size distribution. They did this by moving a vertical line along a scale bar continuously within 3 s. Eleven ticks were evenly spaced over the scale bar to guide participants’ estimation. Initially, the line appeared at a random position over the scale bar with equal probability. In the reproduction task, participants were required to adjust the diameter of a probe ring to match the absolute size of the target ring within 3 s. The physical appearance of the probe ring was identical to the target ring, except for the diameter. The probe ring’s initial size was randomly sampled from the same distribution as the target ring.

In any given block of trials, two tasks were performed in succession as a pair over trials. Depending on which two of the three tasks were performed, there were two types of blocks (Fig. 4a). On each trial of the ‘classification-scaling’ blocks, participants performed the classification task and then the scaling task. On each trial of the ‘classification-reproduction’ blocks, participants performed the classification task and then the reproduction task. For each block type, the granularity level varied between two granularity levels, 2 and 8, on a block-to-block basis. At the start of each trial, a pre-cue (0.5 s) indicated which keys should be used for reporting the class by presenting “45” for granularity level 2 or “12345678” for granularity level 8. Then, a ring stimulus was viewed for 1.5 s and classified by participants within 1.5 s after stimulus onset. After classification, a pre-cue for the scaling or reproduction task, “cv,” appeared for 0.5 s, indicating which keys should be used for scaling and reproduction. Then, a new ring stimulus was viewed for 1.5 s, and participants carried out the scaling or reproduction tasks on that ring. In any given trial, the sizes of the ring for classification and that for scaling or reproduction were independently sampled from the same calibrated (see below) stimulus distribution.

Since any participants in Exp 5 did not take part in Exp 1, we ran the threshold-calibration blocks before administering the training and testing blocks. The threshold-calibration blocks followed the same procedure as Experiment 1, with the exception that the stimuli were solid purple rings (1.219 cd/m2), and two blocks, each consisting of 60 trials, were administered. In the training and testing blocks, we used the purple rings with the same appearance as those in the threshold-calibration blocks. The ring size was randomly sampled for each trial from the distribution calibrated by the threshold-calibration blocks, as in the previous experiments.

As similarly done in Exp 1, the threshold calibration blocks were initially followed by the alternating training and testing blocks for the first 16 blocks (2 blocks for each of the 8 conditions: [testing, training] x [granularity = 2,8] x [reproduction, scaling task]), and then solely by the testing blocks for the remaining 4 blocks (one block for each of the 4 conditions: [granularity = 2,8] x [reproduction, scaling task]). Each testing and training block consisted of 15 and 35 trials, respectively. As a result, a total of 105 testing trials (3 blocks x 35 trials) were acquired for each condition.

The training and testing blocks differed only in that trial-to-trial corrective feedback was provided only in the former. The feedback for the classification task was the same as that of Exp 1. For the scaling task, a green vertical line was presented for 0.5 s at the position corresponding to the true relative size over the scale bar. For the reproduction task, a green ring with the actual size of the target ring was presented for 0.5 s.

### Experiment 6

The procedure for Exp 6 is the same as that of Exp 5, with the following exceptions: (i) stimuli were displayed for 0.3 seconds in both the training and testing blocks; (ii) the ticks were not shown on the scale bar in the scaling task to prevent participants from treating it as a discrete categorization task; (iii) a mask (consisting of white dots randomly distributed around the ring) lasting for 1 s followed the ring stimulus for the scaling and reproduction tasks to minimize aftereffects.

### Experiment 7

In Exp 7, thirty of the individuals who participated in Exp 1 and 2 took part. Exp 7 was conducted to examine whether the granularity can be explained away by the correspondence in motor responses or action space between consecutive trials. The procedure of Exp 7 was the same as that of Exp 3, with the following exceptions. First, the ring size and sound pitch classification tasks were varied on a block-to-block basis. Second, during the pre-cue period, one of the characters, ‘L’ and ‘R’, randomly appeared on each trial to indicate which side of the hands should be used to report. The [1,2,3,4] and [5,6,7,8] keys were assigned to the little, ring, middle, and index fingers of the left hand and he index, middle, ring, and little fingers of the hand, respectively. In the blocks with granularity level 2, [2,3] (or [6,7]) keys were used to report the 1^st^ and the 2^nd^ classes, respectively. In the blocks with granularity level 4, [1,2,3,4] (or [5,6,7,8]) keys were used to report from the 1^st^ to 4^th^ classes, respectively.

There were 36 blocks in total, each consisting of 50 trials. The classification granularity varied between 2 and 4. Exp 7 began with the alternating training and testing blocks for the first 16 blocks (4 blocks for each of the 4 conditions: [testing, training] x [granularity = 2,4]), which were followed by the testing blocks for the remaining 20 blocks (10 blocks for each of the 2 conditions: [granularity = 2,4]).

### Supplementary experiment (Exp S) setup and procedures

Exp S was conducted in an MRI scanner and includes several experimental conditions that differ from the main experiments, as it was optimized for a neuroimaging study. Nevertheless, we analyzed this dataset to examine the impact of ITI on the granularity effect.

Visual stimuli were projected using a projector (Panasonic, PT-DX820) onto a back-projection screen positioned at the end of the magnet bore, with a viewing distance of 100 cm. On the screen, two diagonal gray lines intersected at the center and were continuously displayed in the background to help participants maintain their focus. The stimulus was a gray disk, with its size varying from 0.1437° to 1.4563° in increments of 0.0875°, resulting in a total of 16 stimuli. Choice responses were recorded using an MRI-compatible 8-button response device box. For the current purpose, we included only the experimental sessions where both decision granularity and ITI were manipulated. Each of these sessions included training or testing blocks of trials, which were identical except for feedback and ITI. The training block was at the first block of the session. After that, 3 testing blocks were administered.

In each trial of the testing blocks, participants viewed a pre-cue lasting 0.5 s that indicated the decision granularity of the current trial by displaying the text “45” or “12345678” at the center of the screen. This was followed by a 0.5-s fixation period, a 0.3-s stimulus presentation with a choice response, and a 1.2-s response window. Importantly, the ITI was randomly assigned to be either 2 s or 8 s. In the “45” condition (*G* = 2), participants indicated their choice by pressing one of the buttons labeled “4” and “5” with their left and right index fingers, respectively. In the “12345678” condition (*G* = 8), participants reported their choice using their left and right fingers (excluding the thumbs) for classes “1” to “4” and “5” to “6,” respectively.

The trial structure in the training blocks was the same as in the testing blocks, except that the ITI was fixed at 0.5 seconds, and corrective feedback was displayed for 0.5 seconds after the response window. During the feedback period, the correct choice was shown in red text for each error response, while the color of the fixation diagonals changed from gray to green for each correct response. The number of training and testing trials per block was 65 and 33, respectively, totaling 195 and 297 trials overall.

## QUANTIFICATION AND STATISTICAL ANALYSIS

### Assessing the magnitude of history effects using Kullback-Leibler Divergence (KLD)

*KLD* quantifies the distance between any two given distributions without any parametric assumptions. We utilized *KLD* to measure the impact of each variable on its corresponding choice state. To do so, we quantified the probability distribution of a variable (*s*_*t*_, *c*_*t*−1_, or *s*_*t*−1_) conditioned on each choice state *i* (e.g., *p*(*s*_*t*_|*c*_*t*_ = *i*)). To calculate the probability distribution, first, we counted the number of trials conditioned concurrently on *c*_*t*_, *s*_*t*_, *c*_*t*−1_, or *s*_*t*−1_: *N*(*c*_*t*_, *s*_*t*_, *c*_*t*−1_, *s*_*t*−1_). Then, if we want to calculate the probability distribution of *c*_*t*−1_, we computed 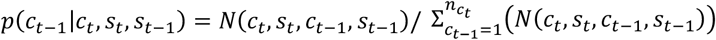. Then, we averaged *p*(*c*_*t*−1_ |*c*_*t*_, *s*_*t*_, *s*_*t*−1_) over *s* and *s*_*t*−1_ :

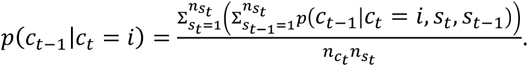

Lastly, *KLD* was determined by taking a *KLD* of *p*(*c*_*t*−1_|*c*_*t*_ = *i*) from the uniform distribution. Consequently, *KLD* reflects the variability of the current choice that can be attributed to the previous choice or stimulus.

### Assessing history effects with a probit regression model with orthogonally decomposed choice regressors

To comprehensively assess the history effects, including the granularity effect, on the current choice, we regressed the current choice *c*_*t*_ onto multiple factors: previous choices (*c*_*t*−*i*_); interactions of previous choice with granularity (*c*_*t*−*i*_ × *G*), with previous RT (*c*_*t*−*i*_ × *RT*_*t*−*i*_), and with current RT (*c*_*t*−*i*_ × *RT*_*t*_); previous stimuli (*s*_*t*−*i*_); interaction of previous stimuli with granularity (*s*_*t*−*i*_ × *G*), with previous RT (*s*_*t*−*i*_ × *RT*_*t*−*i*_), and with current RT (*s*_*t*−*i*_ × *RT*_*t*_); current stimulus (*s*_*t*_); interactions of current stimulus with granularity (*s*_*t*_ × *G*) and with current RT (*s*_*t*_ × *RT*_*t*_).

To ensure a fair comparison of the history effects across varying levels of decision granularity, we developed a regression model with the following features. First, the current choice (*c*_*t*_), as the regressand, was binarized irrespective of the levels of decision granularity: *y*_*t*_ ∈ {0,1}, where 0 represents the classes in the smaller half and 1 represents the classes in the larger half. Second, the previous choice, *c*_*t*−*i*_ was decomposed into three ordered components, 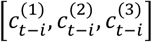 (see Table 1). The first-order component, 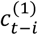, indicates whether *c*_*t*−*i*_ a larger or smaller half. The second-order component, 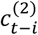, indicates whether *c*_*t*−*i*_ is a larger or smaller half within each of the first-order component. The third-order component, 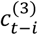, indicates whether *c*_*t*−*i*_ is a larger or smaller half within each of the second-order component. This decomposition allows the first-order component, 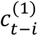, to reflect the influence of the previous choice in a way that is comparable across different granularity conditions. Thus, we used only 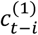 in evaluating the granularity effect. Third, the weighted sum of these ‘orthogonally ordered’ previous-choice components and the other regressors is linked to the regressand (the binarized current choice variable) through the probit function.

The following probit regression model was used for Exp 1 and 2:

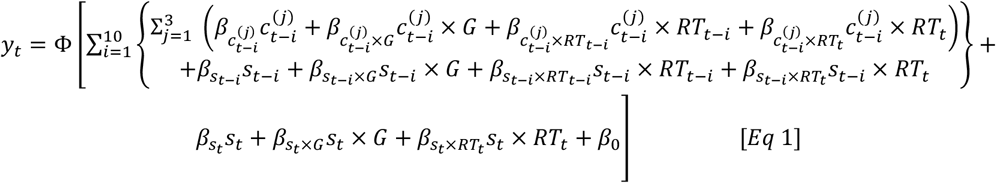

where *β*_0_ accounts for any arbitrary constant bias in choice selection and Φ is the standard cumulative normal distribution. The data from all participants were pooled to calculate the coefficients. The non-interaction regressors such as 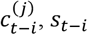, and *RT*_*t*−*i*_ were standardized to have a mean of 0 and a standard deviation of 1 for each block of trials. This block-wise standardization was done to prevent any potential block-to-block history effects from interfering with the trial-to-trial history effects, which are the focus of this study. Consequently, the interaction regressors involving those standardized factors were the products of two standardized factors. The granularity levels, [2, 4, 8], were coded into *G* values, [0, 1, 2]. This allowed 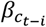 and 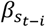 to solely account for the history effects of the previous choice and stimulus, respectively, when the granularity level was 2 by setting the interaction regressor terms involving *G* to 0. Furthermore, the regressor, 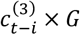, was excluded from the actual regression analysis because 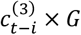 can be reduced to 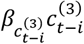. Thus, the regression model included a total of 154 coefficients as the regressors involved previous choices and stimuli up to 10 trials back (*i* = [1,10]). P-values of the probit regression were adjusted to control for false discovery rate (FDR) in multiple hypotheses testing^145^. The total number of hypotheses tested was equal to the number of coefficients, which was 154 for Exp 1 and 2.

To test the robustness of the results on the individual level, we applied the regression model depicted in [*Eq* 1] to the choices of each individual participant. For the robustness of the fitting, we simplified [*Eq* 1] by involving history regressors up to 7 trials back. The statistics of the regression coefficients in this individual level analysis were obtained by conducting t-tests to determine whether participants’ coefficients significantly deviate from 0. The p-values were adjusted to control for FDR, just as we did for the population-pooled data.

The regression model for Exp 3 and 7, where the two tasks alternated on a trial-to-trial basis, was the same as [*Eq* 1], except that the regressors included the previous choices and stimuli only up to 1 trial back (*i* = 1). Consequently, FDR was controlled for a total of 15 hypotheses.

In Exp 4, the regression model was the same as that in Exp 3 and 7, with a few exceptions. The granularity level varied, on a trial-to-trial basis, between 2 and 8, coded into *G* values of 0 and 1, respectively. Hence, *G* had trial indices: *G*_*t*−1_ and *G*_*t*_ when evaluating the granularity effects of the previous and current choices, respectively. Accordingly, for the dataset arranged for evaluating the granularity effect of the previous choice 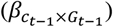, the regression model was as follows:

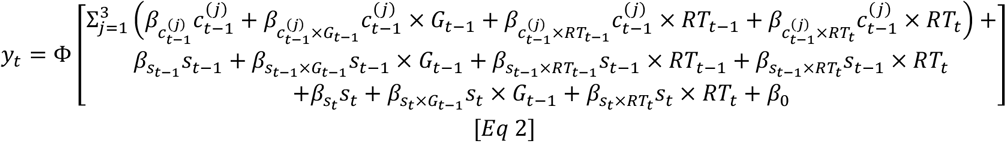

For the dataset arranged for evaluating the granularity effect of the current choice 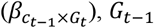 was replaced by [*Eq* 2]. In this case, 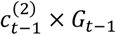 and 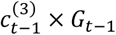 can be reduced to 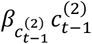 and 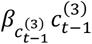, respectively. Thus, these two terms were excluded from the analysis. So, for these regression models, FDR was controlled for a total of 18 hypotheses.

For the reproduction responses on Exp 5 and 6, the regression model was

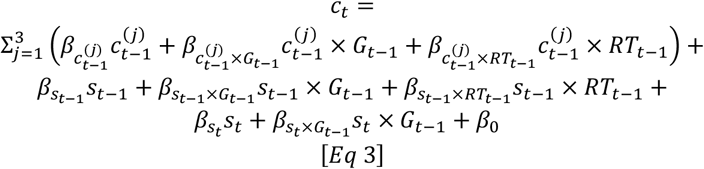

[*Eq* 3] is the same as [*Eq* 2], with the following exceptions. First, there was no link function because the reproduction response is an unbounded continuous variable. Second, the block-wise standardized values of the reproduction responses (*c*_*t*_) were used. Third, there were no regressors involving *RT*_*t*_ because the task performed in the current trial was scaling or reproduction. FDR was controlled for a total of 13 hypotheses.

For the scaling responses on Exp 5 and 6, the regression model was the same as [*Eq* 3], with the following exceptions. First, the probit link function was used because the scaling response is normalized between 0 and 1. Second, the scaling response (*c*_*t*_) was not standardized because its original value is already normalized between 0 and 1.

In Exp S, two regression models were used. One for intuitive visualization of the granularity effect [*Eq* 4], and the other for the rigorous statistical test of the impact of ITI on the granularity effect [*Eq* 5]. The regression model for visualization was

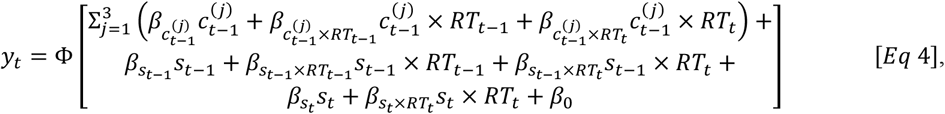

which is the same as [*Eq* 2], except that it does not have the regressors with granularity. This is because we intended to show the raw coefficient of 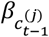 for each granularity level and ITI condition by fitting [*Eq* 4] to all combinations of granularity and ITI. This raw coefficient approach allows for more straightforward interpretations when 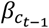 does not significantly deviate from 0 when granularity is 2. If 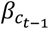 does not significantly deviate from 0 when granularity is 2, it would be unclear whether 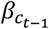 significantly deviates from 0 when granularity is 8 even if 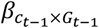 is significantly positive.

The regression model for statistical tests was

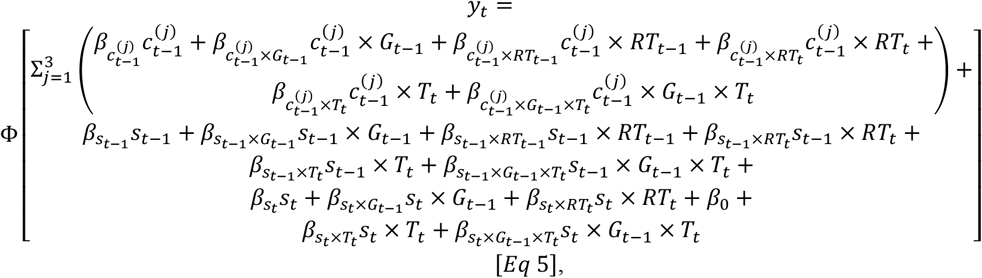

which is the same as [*Eq* 2] except that it includes the interaction terms of ITI (*T*_*t*_). For each of the key regressors, *c*_*t*−1_, *s*_*t*−1_, and *s*_*t*_, the interaction effects of *T*_*t*_ on both key regressors (e.g.,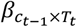) and their granularity effect (e.g.,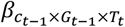) were added to the regression model. Short and long *T*_*t*_ were encoded as -1/2 and 1/2, respectively, for the regression analysis.

### Bayesian model for parallel belief propagation in stimulus and DV spaces

We have developed a Bayesian model to offer a principled and unified account of the history effects observed across the classification, scaling, and reproduction tasks: the stimulus-repulsion bias, choice-attraction bias, and granularity effect (Figure 8A). This model shares many general assumptions with other Bayesian decision-making models^10^, such that: (i) When a task is given, human individuals learn an internal (generative) model of the causal structure of relevant random variables involved in the task, which may or may not reflect the true structure; (ii) As the state of any of those variables is observed or known, they propagate it over the internal model to probabilistically infer the state of the variable critical to the task; (iii) They form a DV and commit to a decision by applying a decision rule to the current state of the DV. In order to explain the history effects within the Bayesian framework, the model further assumes that humans continually update their probabilistic knowledge in their internal model (beliefs) based on their trial-to-trial task episodes (Figure 8D). Specifically, the model assumes that two prior beliefs are updated in parallel, independently of each other, regarding the stimuli and DV, respectively. This assumption of “parallel belief updating in stimulus and DV spaces” prescribes that the stimulus-repulsion bias originates from updating the belief about the stimuli, while the choice-attraction bias and its modulation by decision granularity originate from updating the belief about the DV. The model’s rationale is stated in the main text, while statistical details are provided below.

### Internal model for stimulus inference

The internal model for the stimulus space (*IM*^*s*^) consists of two components of probabilistic knowledge, one about the stochastic causality of external stimuli (*s*) on their sensory measurements (*m*^*s*^; left panel of Figure 8B) and the other about the distribution of external stimuli encountered during the task. They can be expressed by *p*(*m*^*s*^|*s*) = 𝒩(*s, σ*_*s*_) and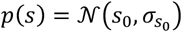, respectively, where 𝒩(*μ, σ*) denotes the normal distribution with mean *μ* and standard deviation *σ*. Observers use the Bayes rule to form a posterior belief about the stimulus *p*(*s*|*m*^*s*^) when the state of *m*^*s*^ is sensed in their sensory apparatus. This is done by propagating the sensed state of *m*^*s*^ over *IM*^*s*^, which is proportional to *p*(*m*^*s*^|*s*)*p*(*s*) (left panel of Figure 8C). As a point estimate of the stimulus ŝ (left panel of Figure 8C), observers take the mode of the posterior belief (maximum a posteriori). Observers adjust the size of the probe ring to align with ŝ in the reproduction task.

### Internal model for DV inference

To perform the scaling and classification tasks, observers need to learn the internal model for the DV space (*IM*^*DV*^). *IM*^*DV*^ also consists of two components of probabilistic knowledge, one about the stochastic causality of DV states (*v*) on their internal measurements (*m*^*v*^; right panel of Figure 8B) and the other about the distribution of DV states formed during the task. Since, *v* unlike *s* in *IM*^*s*^, is bounded between 0 and 1, we defined the shapes of *p*(*m*^*v*^|*v*) and *p*(*v*) by assuming that given a point value of *v*, its measurement *m*^*v*^ diffuses from *v* via Brownian motion while turning its direction at the bounds. We will denote the resultant distribution of the particles of this bounded Brownian motion by *B*(*v, σ*_*v*_), where*v* is the starting point of diffusion and *σ*_*v*_ is the strength of diffusion. *B*(*v, σ*_*v*_) is derived as follows. First, the particles are distributed in an effectively bounded space, [−*b, b*], as a normal distribution, 𝒩(*v, σ*_*v*_). The space is “effectively bounded” because we fixed *b* to 4 while the maximum value of *σ*_*v*_ we have considered in the following simulation was 0.2. Therefore, even if *v* and *σ*_*v*_ have their greatest values 1 and 0.2, respectively, *N*(*v, σ*_*v*_) is effectively 0 at the bound of the space because*N*(*b*; *v* = 1, *σ*_*v*_ = 0.2)/*N*(*v*; *v* = 1, *σ*_*v*_ = 0.2) *<* 10^−48^. Then, 𝒩(*v, σ*_*v*_) defined in [−*b, b*] is truncated into *B*(*v, σ*_*v*_) defined in [0, 1]. In doing so, the part of 𝒩(*v, σ*_*v*_) within [−*b*, 0) is flipped to [0, *b*] and added to the part of 𝒩(*v, σ*_*v*_) within [0, *b*]. We will denote the distribution after this first round of truncation by 𝒩(*v, σ*_*v*_)^1^. On the second round, the part of 𝒩(*v, σ*_*v*_)^1^ within (1, *b*] is flipped to [−*b* + 1,1] and added to the part of 𝒩(*v, σ*_*v*_)^1^ in [−*b* + 1,1]. The distribution after the second round is denoted as 𝒩(*v, σ*_*v*_)^2^. This truncation procedure is repeated *n* times until the part of 𝒩(*v, σ*_*v*_)^*n*^ falling outside of [0,1] is 0. We denote 𝒩(*v, σ*_*v*_)^*n*^ as *B*(*v, σ*_*v*_). Thus, the likelihood of *v* given *m*^*v*^ is defined as *p*(*m*^*v*^|*v*) = *B*(*m*^*v*^, *σ*_*v*_). Then, when the state of *m*^*v*^ is known, a posterior belief about the DV can be formed by propagating the known state of *m*^*v*^ over *IM*^*DV*^: *p*(*v*|*m*^*v*^) *∝ p*(*m*^*v*^|*v*)*p*(*v*). The same estimation rule (MAP) is applied to obtain a point estimate of DV 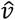 from the posterior distribution. Observers adjust the location of the probe line to align with 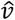 on the scale bar in the scaling task. As for the classification task, observers apply the following decision rule: (i) they establish a set of criteria to evenly divide the DV space based on the decision granularity level ([0.5] for 2; [0.25, 0.5, 0.75] for 4; [0.125, 0.25, …, 0.75,0.875] for 8); (ii) commit to a choice corresponding to a subregion of the DV space within which 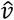 falls.

### Origin of DV measurements

We did not specify the origin of *m*^*v*^ (the evidence for v) when we described *IM*^*DV*^ above, to avoid complications. This specification is essential for connecting the two internal models, *IM*^*s*^ and *IM*^*DV*^, and requires a separate description with a clear rationale and sufficient details. In both scaling and classification tasks, the true state of DV is, by definition, the probability that the current state of the stimulus *s*_*t*_ is larger than the entire possible states of *s*, which is equal to the cumulative distribution function of *s* for the interval from negative infinity to 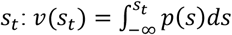. However, *v*(*s*_*t*_) is not directly available since *s*_*t*_ cannot be observed but only inferred from its sensory evidence 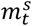. This means that not the true state of DV (*v*(*s*_*t*_)) but only its “evidence” is available. By substituting *s*_*t*_ with ŝ_*t*_, which is inferred from the sensory evidence with the prior knowledge of *s* in *IM*^*s*^, we can define the evidence for 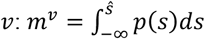 (angled arrow at the top of Figure 8C). To sum in general terms, the outcome of the probabilistic inference based on a sensory measurement with *IM*^*s*^(ŝ) is converted by the absolute-to-relative mapping function (Φ(*s*) = ∫ *p*(*s*)*ds*; the middle panel of Figure 8D) into the evidence for inferring the true state of DV with *IM*^*DV*^(*m*^*v*^). Due to this conversion, the same stimulus estimate ŝ can result in different states of *m*^*v*^ depending on the shape and location of the stimulus prior *p*(*s*).

### Belief updating in stimulus space

So far, we have defined the internal models for the stimulus and DV spaces and described how observers carry out the reproduction, scaling, and classification tasks based on the inferred outcomes with those internal models, ŝ and 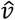. Next, we will specify how some beliefs in the two internal models are updated based on the events occurring over trial episodes. The prior beliefs about stimulus and DV distributions, *p*(*s*) and *p*(*v*), are updated in parallel. Let us denote the stimulus prior used to perform a task on the previous trial by *p*(*s*), the memory of sensory measurement by 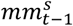, and the stimulus prior that is updated aft should be updated toward er the previous trial and will be used in the current trial by *p*(*s*)′. Then the stimulus prior should be updated toward 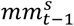 since 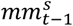 provides new evidence for the stimulus distribution. This updating process is captured by formalizing the prior of *s* in the current trial as the posterior of *s* in the previous trial, as follows: 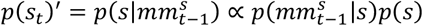 (the left panel of Figure 8D). Here, since 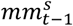 is assumed to be subject to unbiased diffusion error over time in memory, 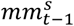 is randomly sampled in each trial from 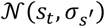, where *s*_*t*_ is the stimulus at trial *t* and 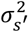 is the sum of sensory and memory variance, 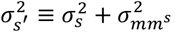. Thus, the likelihood of *s* given 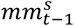 is defined as 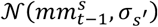.

### Belief updating in DV space

The belief updating process in DV space shares the basic rationale with that in stimulus space: the prior belief about *v* is continually updated over trials. However, whereas the prior belief about *s* is updated by *mm*^*s*^, the prior belief about *v* is updated by decision commitment. Specifically, the state space of *v* is reduced to a subregion corresponding to the states of *v* consistent with the choice. In other words, the prior of *v* for the current trial *p*(*v*_*t*_)^′^ is the probability distribution of *v* conditioned on the previous choice, as follows: *p*(*v*_*t*_)^′^ = *p*(*v*|*choice*_*t*−1_) *∝ p*(*choice*_*t*−1_|*v*)*p*(*v*) (the right panel of Figure 8D). An important implication of this conditioning is that as the decision becomes more granular, the state space of *v* is further reduced. Additionally, the conditioned prior has a memory leak so that the prior can have values greater than 0 outside the range (Figure 1B). This leak on the DV prior was implemented by convoluting it with a normal distribution, 𝒩(0, *σ*_*leak*_). To bound the convolved prior belief within [0, 1], the aforementioned truncation procedure is also applied.

In sum, our Bayesian model account of the history effects in the reproduction, scaling, and classification tasks is defined by the two internal models for inferring the stimulus and DV states (*IM*^*s*^ and *IM*^*DV*^) and the algorithms of belief updating of the prior beliefs in those two internal models, which are fully specified by a total of six free parameters, 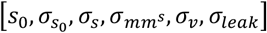.

### Alternative models for the granularity effect

Considering the necessity of updating beliefs in DV space for the granularity effect, we explored previously suggested sources of history effects as potential alternative explanations for the granularity effect identified in our study. To this end, we built three alternative models that implement those suggestions by modifying the original Bayesian model described in the previous section. We emphasize that all of these alternative models share a common feature: none of them assume probabilistic inference or belief updating in DV space. Below, the original Bayesian model will be referred to as ‘the standard model.’

### Sensory adaptation model

The sensory adaptation model assumes that the response gain of sensory neurons encoding the stimulus in the previous trial is reduced. This model is identical to the standard model except for the following. First, the response gain of the sensory neurons tuned to the stimulus measurement in the previous trial 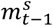 is selectively suppressed, which is incorporated in the model by reducing the gain of likelihood function around 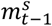. Specifically, the suppressed gain, *f*_*s*_(*x*), is described by 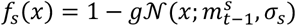, where *g* is the parameter that modulates the strength of gain reduction. Consequently, for a given present stimulus measurement, the likelihood function is modified to 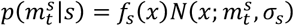. Second, it does not involve any process of belief updating in stimulus space. Third, since there is no informative prior in the DV space, the DV estimate 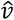 is the same as the DV measurement *m*^*v*^. Four parameters, 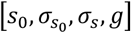, specify the model.

### Stimulus-prior-updating-by-the-stimulus model

The stimulus-prior-updating-by-the-stimulus model is identical to the standard model except that it does not involve any process of probabilistic inference in the DV space. The model shares the same internal model for stimulus inference and the same belief updating in stimulus space as the standard model. Thus, it performs the reproduction task in exactly the same way as the standard model. On the other hand, the model carries out the scaling and classification task using the uninformative DV prior, similar to the sensory adaptation model. However, note that this model does not assume any sensory gain reduction at all, unlike the sensory adaptation model. Four parameters, 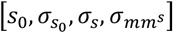, specify the model.

### Stimulus-prior-updating-by-the-choice model

Like the stimulus-prior-updating-by-the-stimulus model, the stimulus-prior-updating-by-the-choice model shares the same internal model for stimulus inference as the standard model. However, this model differs from the stimulus-prior-updating-by-the-stimulus model in that it updates its stimulus prior not only based on the memory of the previous memory 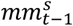 but also based on decision commitment. Specifically, the updated stimulus prior, *p*(*s*)^′^, is updated to be consistent with the previous choice, *c*_*t*−1_, as *p*(*v*)^′^ is updated to be consistent with the previous choice in the standard model. Thus, this model implements the possibility that the granularity might arise in stimulus space instead of DV space. For example, when the level of granularity was 4 and the previous stimulus was classified as the third class, observers consider that the previous stimulus was sampled within the range that is consistent with the third class. This consideration narrows down the stimulus prior to the range corresponding to the third class, which can be calculated based on the stimulus prior to be [Φ(0.5), Φ(0.75)], where Φ is the inverse cumulative distribution of the updated prior, *p*(*s*)^′^. Then, the prior value outside the range is 0. This conditioned prior, *p*(*s*)^′′^, is defined as *p*(*s*)^′′^ *∝ p*(*s*)^′^, where *s* is within [Φ(0.5), Φ(0.75)]. Otherwise, *p*(*s*)^′′^ = 0. Also, it is assumed that the conditioned prior has a leak so that the prior can have values greater than 0 outside the range. This leak is implemented by convoluting a normal distribution, *N*(0, *σ*_*leak*_), with the conditioned prior, *p*(*s*)^′′^. Thus, the model has five parameters, 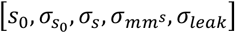.

### DV-prior-updating-by-the-choice model

The DV-prior-updating-by-the-choice model is identical to the standard model except that it does not involve any belief updating in the stimulus space. The model was developed to test whether the DV space belief updating is sufficient to induce not only the granularity effect but also the stimulus repulsion because the skewed DV distribution might have induced the repulsive bias as previous studies suggested^75^. As stimulus updating does not occur, the parameter for stimulus memory diffusion 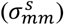 is not required. Thus, five parameters, 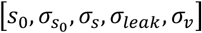, specify the model.

### Model simulation

To determine if the models can qualitatively display the history effects seen in our study’s human participants, we conducted simulations using the following procedure. First, we randomly sampled 1,000 stimulus magnitudes from a normal distribution, 𝒩(0,1). Then, we let the models perform the reproduction, scaling, and classification tasks on this same sequence of stimuli while varying their model parameters. Over these trials, decision granularities were randomly varied trial-to-trial between 2 and 8. Then, the sequence of responds made by the models was regressed onto the regressors involving the stimulus and choice factors, by excluding the regressors of response time from [*Eq* 2] and [*Eq* 3] because the models did not produce response times. To be consistent with the human experiments, we only included the trials in which the level of granularity in the current trial was 2. The entire procedure outlined above was repeated 100 times for each set of parameters, with a newly sampled sequence of 1000 stimulus magnitudes for each repetition. Then, the regression coefficients acquired from these 100 repetitions were averaged.

The model parameters used in the above simulations were determined as follows. To thoroughly and impartially explore the parameter space, we used all combinations of parameters selected from 10 evenly spaced parameter values. The specified range of the parameter space can be found in Table 2. For simplicity, we set the mean of the stimulus prior, *s*_0_, as 0 for all models because the overall prior bias would not induce systematic bias to history effects. As a result, there were 10^3^ parameter sets for the sensory adaptation and stimulus-prior-updating-by-the-stimulus models, 10^4^ for the stimulus-prior-updating-by-the-choice model, and 10^5^ for the standard (original Bayesian) model.

**Table 2.** The range of parameters used in model simulation. For each parameter, 10 values were selected for the model simulation, evenly spaced within the given parameter range. For example, if the range is [1, 10], the parameter values tested are the integers from 1 to 10.

|  | Sensory adaptation model | Stimulus-prior-updating-by-the-stimulus model | Stimulus-prior-updating-by-the-choice model | DV-prior-updating-by-the-choice model | Standard model |
| --- | --- | --- | --- | --- | --- |
| $\sigma_{s_0}$ | [0.4, 4] | [0.4, 4] | [0.4, 4] | [0.4, 4] | [0.4, 4] |
| $\sigma_s$ | [0.1, 1] | [0.1, 1] | [0.1, 1] | [0.1, 1] | [0.1, 1] |
| $g$ | [0.1, 1] | - | - | - | - |
| $\sigma_{mm^s}$ | - | [0.4, 4] | [0.4, 4] | - | [0.4, 4] |
| $\sigma_{leak}$ | - | - | [0.2, 2] | [0.02, 0.2] | [0.02, 0.2] |
| $\sigma_v$ | - | - | - | [0.02, 0.2] | [0.02, 0.2] |

From these simulation results, we selected the best simulation parameter set as the one with the smallest sum of errors in the regression coefficients compared to human participants in Exp 4, 5, and 6 as follows. First, we excluded the parameters that generated classification accuracies (e.g., correct choice proportion) outside of 95% CI of human subjects in Exp 4. Then, we calculated the sum of absolute errors of 8 coefficients between the model and that of scaling and reproductions reports 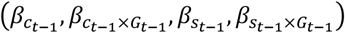 combining data of Exp 5 and 6 (Figure 8G).

## SUPPLEMENTAL INFORMATION

Document S1. Figure S1.

**Figure S1.**
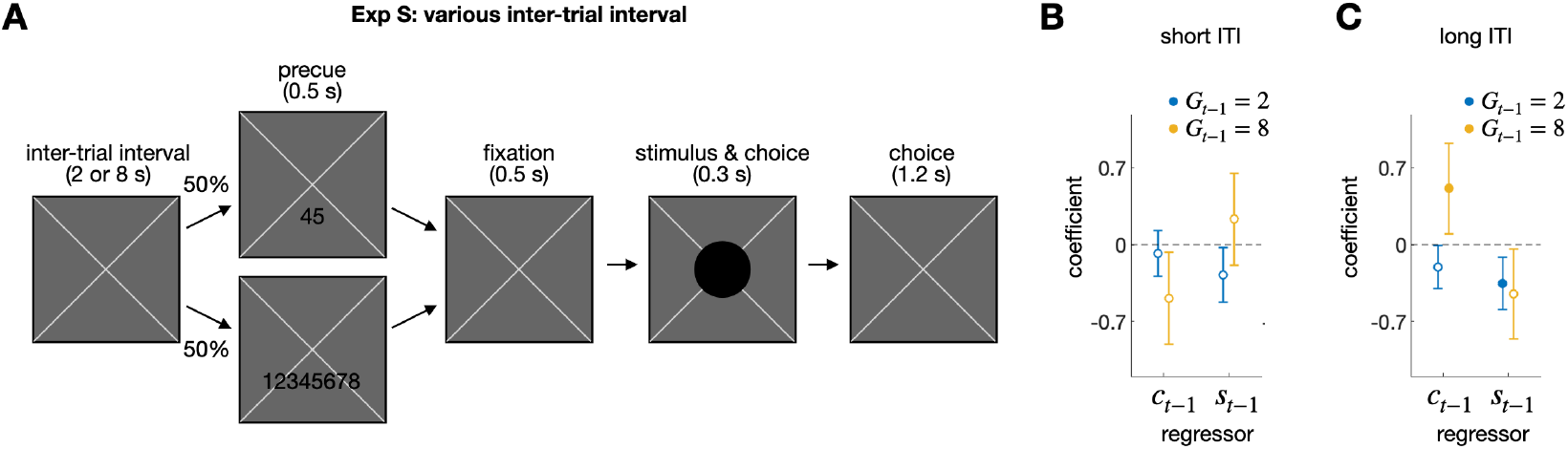
Supplementary experiment (Exp S) with varying durations of ITI, related to STAR Methods. (A) Trial structure. The inter-trial interval (ITI) was randomly assigned either a short (2 s) or long (8 s) duration for each trial. Participants classified the size of a disk into various categories. Data were collected while scanning participants’ brain activity inside a 7T MRI scanner. Unlike in the main experiments (Exp 1-7), the threshold for size contrast was not calibrated individually; instead, one fixed distribution was employed for all participants. This led to a size distribution that was significantly wider than the one used in the main experiments, occasionally showing extremely small or large sizes. To align the task difficulty of Exp S with that of the main experiments, we excluded stimuli from the analysis that significantly deviated from the mean, corresponding to the extreme 28 % of the original distribution. (B,C) Regression of the current choice onto the previous choice (*c*_*t*−1_) and stimulus (*s*_*t*−1_) for the short (B) and long (C) ITI conditions. The regression coefficients are shown separately for decision granularities of 2 (blue circles) and 8 (yellow circles). The filled circles indicate that the corresponding coefficients significantly deviated from 0 after being controlled for FDR.

